# MYH9, a cytosolic myosin protein, binds to dengue virus 3’UTR and facilitates replication and cellular entry

**DOI:** 10.64898/2026.03.02.708980

**Authors:** Deepti Maisnam, Deepika Rathore, Lekha Gandhi, Preeti Chauhan, Musturi Venkataramana

**Affiliations:** Department of Biotechnology and Bioinformatics, School of Life Sciences, University of Hyderabad, Prof. C. R. Rao Road, Gachibowli-500046, Telangana, India

**Author notes:** Corresponding address: Email id.

**Keywords:** Dengue virus, MYH9, Virus receptors, Severe dengue, Flaviviruses, UTRs

## Abstract

Dengue infections are considered an increasing threat to mankind due to their rapid global spread rate. The development of a widely accepted drug/vaccine is hindered due to an incomplete understanding of the virus lifecycle. Present data suggest that a cytoskeleton protein, called MYH9 binds to the 3’UTR, at A4 region, a highly conserved part of the UTR across the serotypes. The levels of this protein were found to be elevated in the cells infected with the virus and the above increase is commensurate with the virus load. This protein is found to accumulate at the endoplasmic reticulum (site of virus replication) and interacts with dsRNA (a replicative intermediate), suggesting its involvement in replication. Inhibition of this protein’s expression by its siRNA reduced viral load, supporting its role in viral replication. Immunofluorescence studies indicate that this protein accumulates at the cell periphery and pulldown studies suggest that this protein interacts with the viral envelope protein, suggesting a role in the dengue virus’s cellular entry, possibly by acting as a receptor. Use of an anti-MYH9 drug, ML-7 indicated the reduction of the virus load, prevented the accumulation at the periphery and aided in regaining the cell morphology of virus infected cells, confirming its role in replication and entry. Collectively, these studies demonstrate a dual function of MHY9 in the virus life cycle, which may serve as a general paradigm for the other viruses and hence to develop specific drugs.

## Introduction

Dengue, one of the mosquito borne viral diseases, causes nearly 400 million infections annually [1]. The year 2023 has been reported to have recorded the highest number, with 6.5 million cases. This increase has caused fear in the general public and led to an alarming situation globally. Dengue infections caused by four serotypes DENV1-4 created havoc in the tropical and sub-tropical regions, with disease symptoms ranging from asymptomatic conditions to symptoms leading to severe cases, sometimes to death. The symptoms of this virus infections are categorised by WHO, as with and without warning signs [1]. These symptoms range from mild to severe, including fever, headache, rashes and body pain. In a few cases, it may lead to complications like dengue haemorrhagic fever (DHF) or dengue shock syndrome (DSS). Factors like dengue virus serotype variation, antibody dependent enhancement (ADE), variation in immune response, antibodies against NS1, and autoimmunity are reported to contribute to the severity [2]. This virus belongs to the genus *Orthoflavivirus* of the virus family *Flaviviridae*. The genome of this virus is a positive sense RNA with 10.7 kb size, codes for the polyprotein that is composed of three structural (capsid, pre-membrane and envelope) and seven non-structural proteins (NS1, NS2A, NS2B, NS3, NS4A & B, NS5) along with two untranslated regions (5’ and 3’UTRs) flanking the genome. These proteins and the UTRs play an indispensable role in the viral life cycle, from entry into the cells through replication, translation and progeny virus production [3]. While residing in the cytoplasm and playing a critical role in the virus’s life cycle during infection, a few of these proteins localise to different subcellular compartments, thereby mitigating their functional activities. In this direction, capsid protein is reported to be localised to the nucleus, NS3 to the nucleus and mitochondria and triggers mitochondrial dysfunction [4,5]. This kind of downplaying host resistance by different proteins is common in many viruses [6,7]. Besides the viral proteins, the virus hijacks several host factors to survive within host cells. Interactions between the virus and complex host factors represent an important step in studying viral pathogenesis. In this direction, the interactions and the role of host proteins like poly ‘A’ binding protein (PABP), poly tract binding protein (PTB), human lupus antigen (La) with virus components of hepatitis C virus (HCV), coxsackie virus B (CVB), foot and mouth disease virus (FMDV) and dengue virus are well established [8]. Efforts are being made to understand the multiplexed nature of viral pathogenesis and the interacting proteins or mechanisms of interaction are being targeted to develop interventions [9]. Understanding such interactions and mechanisms poses challenges, as they are unique to different viruses and complicate. However, to develop therapeutics, it is pivotal to understand such interactions, which can eventually be explored as drug targets [10].

In case of dengue virus infections, all ten proteins and two untranslated regions are reported **to** interact with different host derived proteins [11]. A few of these proteins are known to assist in the multiplication of the dengue virus at different stages and to enhance pathogenesis [12–14]. In this study, MYH9 a cytoskeleton protein, was found to interact directly with the dengue virus 3’UTR and was found to be involved in virus replication and entry. The levels of this protein were found to be elevated during the infections and in turn, the virus load was also enhanced. This protein was found to be involved in two important steps of the virus life cycle, replication and entry. The study also shows that this interaction is a potential drug target, as analysed by siRNA and the drug ML-7 targeting MYH9.

## Materials and methods

### Cell lines and virus cultures

Vero-E6, Baby hamster kidney (BHK) Human embryonic kidney (HEK293), Hepatocyte carcinoma G2 (HepG2), and Human erythroleukemic K562 cell lines were purchased from National Centre for Cell Science (NCCS), Pune, India, maintained in Dulbecco’s modified Eagle’s medium (DMEM), [K562 in RPMI + L-Glutamine, (Gibco)], at 37℃ and 5% CO_2_. The media were supplemented with 10% Foetal Bovine Serum (FBS) and 1% antibiotic-antimycotic solution (anti-anti, Gibco). The virus cultures, confirmation and quantification were performed as described earlier from our lab [5]. DENV 1&2 strains used in the present study for cell culture infections were isolated from strains circulating in Hyderabad, India in 2019, propagated in Vero-E6 cells, quantified and further characterised [15]. The viral supernatants were diluted in DMEM and stored at -80℃ for further experiments.

### Antibodies

The sources and used dilutions of the antibodies: Polyclonal anti-MYH9 (Proteintech, 1:10000), anti-actin (Santa Cruz, 1:1000), anti-GAPDH (Proteintech, 1:10000), anti-NS2BNS3 (in-house raised, 1:500), anti-calnexin (Merck, 1:200), anti-dsRNA (Merck, 1:200). Horse radish peroxidase (HRP)-conjugated rabbit anti-mouse and mouse anti-rabbit IgG are from GeNei (Bangalore, India: 1:10000), anti-rabbit IgG (H+L) CF^TM^488A secondary antibody (Merck), goat anti-rabbit IgG antibody (Dylight594, Genetex, 1:1000) and anti-mouse IgG Fab2 Alexa Fluor (R) 594 (Cell Signalling Technology, 1:1000).

### Construction of recombinant plasmids

The DENV1 5’ and 3’ UTRs were amplified by PCR using the complete genome clones (Acc No. KX618705, characterised in our lab) as templates. The amplification reaction was performed in a final volume of 25 μL containing 2.5 mM each of four dNTPs, 1x PCR reaction buffer, 0.5 U of DNA polymerase, 100 ng of template and 10 pM of each forward and reverse primer. The primer sequences for 5’ UTR forward 5’CCG<u>GAATTC</u>TAATACGACTCACTATAGGGTTAGTCTACGTGGAC3’ and reverse 5’ <u>TCTAGA</u>CTCTCGCGTTTCAGCATATTG3’. For 3’ UTR forward 5’CCG<u>GAATTC</u>TAATACGACTCACTATAGGGTGGTAAGCCAACACACTC3’ and reverse 5’ CT<u>GCGGCCGC</u>AGAACCTGTTGATTCAA3’. The PCR amplification was carried out with an initial denaturation step at 95°C for 4 min, followed by 35 cycles of 94°C for 30 sec, 55°C for 30 sec and 72°C for 1 min; and a final extension at 72°C for 10 min. Both 5’ and 3’ UTR amplified products were purified by Thermo Fisher Scientific gel extraction kit method. The gel purified DNA products of 5’UTR and pBluescript KS(+) vector were subjected to double digestion with *EcoRI* and *XbaI* restriction enzymes in a 25 μL reaction at 37^0^C for 3 hrs. The reaction mixture contains 1 μg insert or pBluescript KS(+) vector, 1x cut smart buffer, *EcoRI* 5 U, *XbaI* 5 U, final volume made up of with autoclaved water. The digested products were resolved on 1.2 % agarose gel electrophoresis, and the digested products were excised and gel purified. Similarly, the amplified 3’ UTR DNA and pBSKS (+) were subjected to double digestion with *EcoRI* and *Not I* restriction enzymes and processed by following the above mentioned protocol. The double digested 5’ and 3’ UTR fragments were ligated to linearised pBSKS (+) vector by using T4 DNA Ligase. The reaction contains a final volume of 20 μL with 100 ng of vector, 3-fold excess of the insert, 1 μL of T4 DNA Ligase and 1x T4 DNA ligase buffer. The ligation reaction was performed either at 4°C overnight or at 22°C for 2 hours, transformed and the recombinant clones were confirmed by restriction digestion.

In order to prepare the biotinylated RNA transcript, the recombinant pBSKS (+) vectors containing DENV 5’/3’UTRS were linearized using *Xba I/Not I* (NEB) restriction enzyme at 37°C for overnight and gel excised. Biotinylated RNA transcripts were generated by using the linearised plasmid as a template and MEGAscript T7 kit as per the manufacturer’s instructions. Briefly, 20 µL of transcription mixture containing 1X transcription buffer, 1.25 µL of 20 mM biotinylated UTP and 20U T7 RNA polymerase was incubated at 37°C for 2 hours. Then, 1 µL TURBO DNase was added to the reaction mix and incubated for 15 min at 37°C. 30 µL of nuclease-free water and LiCl precipitation solution were added to the reaction mix and incubated overnight at –20°C or –70°C to precipitate the RNA. The RNA was pelleted by centrifugation at 14,000 rpm for 15 min at 4°C. The pellet was washed with 1 mL of 70% ethanol prepared with DEPC-treated water and re-centrifuged to remove unincorporated nucleotides from the transcription reaction. Then, the RNA was resuspended in 50 µL of nuclease-free water, quantified by Nanodrop spectrophotometer and stored at –20°C or –70°C.

To prepare cell lysates, the thawed cell suspensions of BHK, HepG2, and HEK 293T were transferred into 3 mL of pre-warmed complete DMEM containing 10% (v/v) FBS. The cells were washed twice by centrifugation at 1500 rpm for 3 minutes. 5 mL of complete DMEM medium was added to the pellet, cells were resuspended and transferred to the T-25 culture flasks. The cells were maintained at 37^°^C under 5% CO_2_ until cell confluency reached ∼80%. Then the cells were harvested, washed three times with cold 1XPBS, resuspended in CHAPS buffer (10 mM Tris–HCl pH 7.4, 1 mM MgCl_2_, 1 mM EGTA, 0.5% CHAPS, 10% glycerol, 0.1 mM PMSF, 5 mM 2-ME), and incubated on ice for 30 min for swelling and to lyse the cells. Cell lysates were subjected to centrifugation at 14,0000 rpm for 10 min at 4°C and the supernatants were collected. Protein concentration was determined using the Bradford reagent (Bio-Rad) with bovine serum albumin (BSA) as a standard. The lysates were also prepared using the RIPA (50mM NaCl, 0.5% sodium deoxycholate, 0.1% SDS, 50mM Tris pH 8.0, 1% NP-40) buffer containing protease inhibitor.

### Pulldown assay

A total reaction volume of 100 µL containing 100 µg of cell lysate (approx. 10µL), 12.5 pmol of biotinylated 5′ or 3′ UTR RNA (5 µL), and 85 µL of RNA mobility shift buffer (5 mM HEPES pH 7.1, 40 mM KCl, 0.1 mM EDTA, 2 mM MgCl_2_, 2 mM dithiothreitol), 1unit RNasin and 0.25 mg/ml heparin was incubated at 30°C for 15 min. Then, 200 μl of streptavidin MagneSphere paramagnetic particles (Promega) were added to the reaction and incubated for 10 min at room temperature. The complexes were then washed with the RNA mobility shift buffer five times, without heparin. 30 μl of 2X loading buffer was added to the beads and the mix was incubated at RT for 10 min to dissociate the RNA-protein complexes. The samples containing bound proteins were boiled and loaded onto 10% SDS–PAGE, visualised by silver staining. The interested protein bands were excised from the gel and analysed by MALDI-TOF.

The protein identified by MALDI-TOF was further validated by western blotting. After performing a similar RNA pulldown assay, the flowthrough, washes, and eluates were mixed with 2X loading buffer and heated at 95 °C for 5 minutes to denature, then resolved on 10% SDS-PAGE. The separated proteins were transferred onto a PVDF membrane and blocked with 5% nonfat milk in PBST. The membrane was then probed with anti-MYH9 primary antibody overnight at 4°C. After washing, the membrane was incubated with HRP-conjugated anti-rabbit secondary antibody for 1 hour at room temperature. The protein bands were detected using chemiluminescent substrate and visualised with an imaging system.

### RNA immunoprecipitation

Approximately 1×10⁶ A549 cells/dish were seeded in 100 mm culture dishes. After adherence, the cells were infected with DENV1. Following a 2-hour adsorption period, the medium was replaced with fresh DMEM supplemented with 2% FBS, and cells were incubated for 24–48 hours to allow viral multiplication. Then, the cells were cross-linked with 1% formaldehyde to stabilize RNA-protein interactions, and lysed using RIPA buffer. The lysates were sonicated to fragment nucleic acid-protein complexes and incubated with Protein A/G magnetic beads for 2 hours at 4 °C. The lysate was subjected to centrifugation at 14, 000 rpm for 15 min at 4 °C. The supernatant was collected and incubated overnight at 4°C with anti-MYH9 antibody to capture MYH9-associated complexes. Then, fresh Protein A/G beads were added and incubated for 1 hour at 4°C to pulldown the immunocomplexes. The immunoprecipitated complexes were washed thrice with RIPA buffer. 750 µL of TRIzol was added to the bead-bound complexes and mixed vigorously to ensure complete dissolution. After a 5-minute incubation at RT, 250 µL of chloroform was added, and the tubes were inverted to mix. The samples were centrifuged at 14,000 rpm for 15–20 minutes at 4°C, and the upper aqueous phase was transferred to fresh DEPC-treated tubes. An equal volume of chilled isopropanol was added, incubated overnight at −80°C to precipitate RNA. Centrifugation was carried at 14,000 rpm for 15–20 minutes 4°C to pellet the RNA. Pellets were washed twice with 70% ethanol (centrifuged at 14,000 rpm for 10 minutes each), air-dried, and resuspended in 20 µL of RNase-free water. RNA concentration and purity were assessed using spectrophotometry. Using 1µg of the above isolated RNA as template, RT-PCR was performed with dengue virus 3’UTR reverse primer (5’TGCCTGGAATGATGCTGTAGA’3). The obtained cDNA was further amplified with PCR master mix (GCC biotech) using both forward (GGTAAGCCAACACACTCATGAAA) and the above reverse primer. The PCR conditions: initial denaturation at 95 °C for 5 min, followed by 95 °C for 1 min, 55 °C for 30 sec, 72 °C for 30 sec, 30 cycles of the above three steps and final extension at 72 °C for 5 minutes. The amplicons were analysed by 1% agarose gel electrophoresis and recorded.

### *In-silico* prediction of 3’UTR-MYH9 interactions

Using the RPIseq webserver (http://pridb.gdcb.iastate.edu/RPISeq/), the binding site of the MYH9 protein on dengue virus 3’UTR was determined. The protein sequence and different regions of the 3’UTR sequences (Variable, A2, Variable + A2, A3, Variable + A3 and A4 regions) were used. Regions with highest Random Forest (RF) and Support Vector Machine (SVM) scores was selected as probable binding site.

Further, to determine whether the MYH9 protein binds to all DENV serotypes, 3’UTR sequences of DENV1-4 were obtained from the GenBank database with the accession numbers MG560149, KT282380, ON123662, and KX845005, respectively. The A4 segments of all four serotypes were manually extracted from each sequence based on their nucleotide coordinates and aligned using Clustal W in BioEdit software.

### Levels of MYH9 by western blot analysis

To analyse MYH9 levels during infection, a western blot was performed on Vero-E6 cell extracts infected with DENV1 and DENV2 separately for 72 hours. 20µg of total protein of each sample was loaded onto the 10% SDS-PAGE and transferred onto PVDF membrane. The blot was blocked with skimmed milk solution and probed with anti-MYH9 primary, followed by HRP conjugated mouse anti-rabbit secondary antibodies. The protein bands were detected using chemiluminescent substrate and visualised with an imaging system. The membrane was stripped and reused to probe with anti-NS2BNS3 and anti-actin antibodies as primary antibodies, separately followed by HRP conjugated rabbit anti-mouse and HRP conjugated mouse anti-rabbit secondary antibodies.

### Levels of MYH9 by immunofluorescence assay

An Immunofluorescence assay was performed using Vero-E6 cells infected with DENV1 for 72 hours. Then, the cells were fixed with 4% paraformaldehyde for 20 minutes, followed by permeabilisation using 0.1 % Triton X-100. Further, 2% BSA was used to block the cells, followed by incubation with anti-MYH9 primary and anti-rabbit IgG (H+L) CF^TM^488A secondary antibodies at room temperature. The nuclei were stained with 1X Hoescht stain and analysed under Zeiss Fluorescence microscope at 60x magnification. The supernatants of the above experiment were used to isoalte the virus RNA and to perform the RT-PCR with dengue NS5 primers.

To analyse the MYH9 levels in dengue virus infected clinical samples, the sample collection and processing were carried as described in our earlier studies [5]. Approximately 200 µL of blood sample was collected from the patients infected with dengue virus. Since serum contains ample amounts of albumin, the samples were subjected to albumin out [16]. The protein concentration was estimated by Bradford assay. 20 µg of serum protein was loaded onto 10% SDS-PAGE, resolved and transferred onto the PVDF membrane. The membrane was probed with MYH9 primary antibodies and processed further.

### Overexpression of MYH9 to check viral load

The recombinant plasmid pMYH9-WT-V5 was purchased from Addgene, USA (Plasmid #183512). K562 cells were transfected with 4µg of above plasmid using Lipofectamine 2000 at 1:3 ratios and incubated for 48 hours. Then, the cells were infected with DENV1 and incubated for another 14-16 hours. Total cell extract was made and an equal amount of protein was loaded onto 10% SDS-PAGE along with 3x loading dye. The resolved proteins were transferred onto PVDF membrane, blocked and probed with anti-NS2BNS3 primary antibodies followed by HRP-conjugated mouse anti-rabbit secondary antibodies. The protein bands were developed using chemiluminescent substrate and visualized using an imaging system. The membrane was stripped and reused for GAPDH primary antibodies. In order to confirm the expression of MYH9 in the above experiment, western blot was performed using anti-MYH9 primary antibodies, followed by HRP-conjugated mouse anti-rabbit secondary antibodies. The protein bands were developed as above. The membrane was stripped and reused for GAPDH primary antibodies. RT-PCR amplification was performed from the culture supernatants of the above experiment using primers (Forward 5’CAATATGCTGAAACGCGAGAGAAA3’ and Reverse 5’ CCCCATCTATTCAGAATCCCTGCT3’) for 170 bp at the 5’ end of the virus genome.

### MYH9 and Calnexin double immunofluorescence analysis

Double immunofluorescence staining was performed to examine colocalization of MYH9 and calnexin in DENV-1–infected cells. A549 cells were infected with DENV-1 and incubated for 12–14 h under standard culture conditions. Following incubation, cells were washed with phosphate-buffered saline (PBS), fixed with 4% paraformaldehyde, and permeabilized using 0.1% Triton X-100. Cells were then incubated with primary antibodies against calnexin (mouse; overnight at 4 °C) and MYH9 (rabbit; 2 h at room temperature). Subsequently, cells were washed with PBS and incubated sequentially with species-specific secondary antibodies for 1 h each: anti-rabbit IgG (H+L) F(ab’)₂ conjugated with CF488A for detection of MYH9, and anti-mouse IgG Fab₂ Alexa Fluor® 594 for detection of calnexin. Nuclei were counterstained with 4′,6-diamidino-2-phenylindole (DAPI) for 10 min. After final washes, stained cells were mounted and visualized under a Zeiss fluorescence microscope at 60× magnification. Images were acquired using identical exposure settings for all experimental groups.

### Co-localization analysis for dengue virus dsRNA and MYH9 association

Colocalization analysis was performed to evaluate the spatial association between MYH9 and viral replication intermediates in DENV-1–infected cells. A549 cells were infected and incubated for 12–14 h under standard culture conditions. Cells were subsequently washed with phosphate-buffered saline (PBS), fixed with paraformaldehyde, and permeabilized using Triton X-100. Following permeabilization, cells were incubated with primary antibodies against double-stranded RNA (mouse; overnight at 4 °C) followed by MYH9 (rabbit; 2 h at room temperature). After washing with PBS, cells were incubated for 1 hr with respective fluorophore-conjugated secondary antibodies ie anti-rabbit IgG (H+L) CF™488A to detect MYH9 and anti-mouse Alexa Fluor® 594 to detect dsRNA. Nuclei were counterstained with 1× Hoechst solution, followed by final washes and mounting. Fluorescence images were acquired under a Zeiss fluorescence microscope at 60× magnification.

### Inhibition of DENV1 infection by siRNA of *MYH9* mRNA

siRNAs for *MYH9* mRNA and scrambled RNA [(scRNA) (negative control)] were designed and synthesized (Gene X, Hyderabad, India). The sequence of the siRNA was 5’GCACAGAGCUGGCCGACAAUU3’ and scRNA was 5’ GUGGACAACGGGCACUACCAU 3’. Initial titration experiment was carried to find the optimum siRNA concentration. For this purpose three different concentrations (50, 100 and 200 nm) of siRNA was trasfected using the Lipofectamine 2000 and incubated for 12 hrs. The cell extract was made and the total protein was resolved on 10% SDS PAGE, transferred onto PVDF membrane and probed with anti-MYH9 primary antibodies follwed by HRP lablled secondary antibodies. Them siRNAs were transfected using Lipofectamine 2000, followed by infection with DENV1. The cells were harvested along with the supernatants after 12-14 hrs and the levels of viral protein NS2BNS3 were determined by usibg anti-NS2BNS3 primary antibodies in western blotting. Further, RT-PCR was carried using the isolated virus RNA as template with the primers for 170 bp at the 5’ end of the virus genome.

### MYH9 cellular localisation study

Cellular localisation of the MYH9 protein under virus infection was carried as described earlier [17,18]. For this purpose, Vero E6 and K562 cells were infected with DENV1 and incubated at 4℃ for 2 hours to allow virus adsorption. Then, the cells were shifted to 37 ℃ for virus entry analysis. At 2 and 5 minute time points, the cells were fixed with 4% paraformaldehyde for 15 minutes at RT and then incubated with anti-MYH9, followed by anti-rabbit IgG (H+L) CF^TM^488A antibodies. The nuclei were stained with 1xHoechst stain, observed under Zeiss Fluorescence microscope at 60x magnification and the images were recorded.

### Interactions of MYH9 and DENV1 envelope protein

The crystal structures of dengue virus envelope protein (PDB ID: 4UTC) and MYH9 (PDB ID: 4ZME) were retrieved from the RCSB Protein Data Bank. Protein-protein docking was carried out using the ClusPro server, which employs a fast Fourier transform (FFT)-based algorithm for rigid-body docking of macromolecular complexes. The lowest energy docked model was visualised using PyMOL software (PyMOL Molecular Graphics System Version 3.0.3). The docking pose and the interaction interface between the envelope protein and MYH9 were analysed for potential interacting spots.

Further, the pulldown assay was carried out using purified dengue virus envelope protein and K562 cell lysate. For this purpose, 500 µg of extract was initially incubated with Ni-NTA beads, then 200 µg purified envelope protein was added. The flowthrough, washes and elutes were collected and subjected to 10% SDS-PAGE. The gels were stained with Coomassie and observed for protein bands in the experimental samples. The western blot was performed on the above eluates with anti-MYH9 primary antibodies for further validation. K562 extract alone was processed as a negative control.

### Inhibition of DENV1 infection by anti-MHY9 drug ML-7

Vero-E6 cells were infected with DENV1 and treated with different concentrations (5, 10 and 15 µM) of the anti-MYH9 drug ML-7. The cells were then probed with anti-NS2BNS3 primary antibodies and subsequently with goat anti-rabbit IgG antibody [Dylight 594]. The nucleus was stained with 1X Hoechst stain. The slides were observed under a fluorescence microscope to detect the viral protein. After acquiring the images, the results were analysed by Image J software, and respective graphs were plotted as mean fluorescence intensities of individual cells. From the culture supernatant of the above experiment, viral RNA was isolated using QiAamp viral RNA isolation kit as per the manufacturer’s instructions, followed by RT-PCR amplification using dengue virus specific primers.

### Inhibition of the cellular localisation of MYH9 by ML-7

K562 cells were pre-treated with 15 µM of ML-7 for 1 hour, washed with 1X PBS and then infected with DENV1. After incubation at 4 ℃ for 2 hours the cells were shifted to 37 ℃ for 2 and 5 minutes. Then the cells were removed and processed as previously described to analyse the cellular localisation of MYH9 during virus infection and in the presence of the drug ML-7.

### Cell morphology analysis after ML-7 treatment

For cell morphology analysis, Vero-E6 cells were seeded in a 6 well plate at a density of 1x 10^5^ cells per well. The cells were pre-treated with 5, 10 and 15 µM concentrations of ML-7 and infected with DENV1 strains. After allowing the virus to adsorb for 2 hours, the cells were supplemented with DMEM containing 2% FBS and 1% anti-anti solution. The cell morphology was monitored after every 24 hours and images were captured under a microscope.

### Statistical analysis

The graphs were presented using GraphPad Prism 10. Experiments were performed at least 3 times, and data was represented as mean and standard deviations were statistically analysed by Student’s t-test and ONE-WAY ANOVA. (*), (**) signifies p-values of < 0.05 and < 0.005 respectively. ImageJ software was used to analyse the fluorescence microscope images.

## Results

### MYH9 is a DENV 3’ UTR binding protein

To identify *trans* acting factors that bind to the UTRs of dengue virus, an RNA pulldown assay was performed using an *in vitro* transcribed biotin labelled DENV1 5’ and 3’UTR RNAs and lysates of BHK, HepG2, and HEK cell lines. The RNA pulled down proteins were resolved on 10% SDS PAGE and visualised by silver staining [Figure 1(A)-(B)]. Although there were multiple protein bands that interacted with both UTRs, we focused specifically on the proteins that interacted with 3’UTR, as this UTR is known to play a critical role in genome replication. We selected four specific bands that were absent in the 5’UTR lanes. The molecular weights of the selected bands were approximately 230, 200, 170, and 40 kDa [Figure 1(B)]. The protein bands were gel eluted and subjected to MALDI-TOF analysis. The proteins in the bands were found to be the 1. non-muscle myosin heavy chain II A (also termed MYH9, 230 kDa), 2. AT8B4 –Probable phospholipid-transporting ATPase IM (200 kDa), 3. Ephrin type-A receptor 3 (170 kDa) and 4. Ribosomal protein S2 (40kDa) [Figure S1, A & B). Upon literature analysis of the above proteins, MYH9 was found to be interesting as it was reported to be involved in the life cycle of other viruses also [17–21]. Hence, the elutes of the above pulldown experiment were further subjected to western blot analysis using anti-MYH9 antibodies and the data showed that the eluted band with MW 230 kDa was indeed the MYH9 protein [Figure 1, C – E]. Further, RNA immunoprecipitation (RIP) was performed to confirm the interaction between the 3’UTR and MYH9 under the infected condition. RT-PCR was carried using the obtained immunoprecipitated RNA as template and 3’ UTR specific primers. The results showed an amplicon of 450bp in both DENV 1&2 infected samples [Figure 1, F&G)]. This observation confirms the interaction between dengue virus 3’UTR and MYH9. The binding region of the protein within the 3 ′UTR was predicted using the *in-silico* tool RPISeq, the RF and SVM classifiers yielded probability scores of 0.6 and 0.8, respectively, for the A4 region, indicating a potential interacting region [Figure S2]. In order to confirm that MYH9 interacts with 3 ’UTRs of all four serotypes, we have aligned the A4 regions and the data suggested a high % of homology (92-96%) across the serotypes, suggesting the high possibility of MYH9 binding to the UTRs of all serotypes [Figs S3-S5].

**Figure 1:**
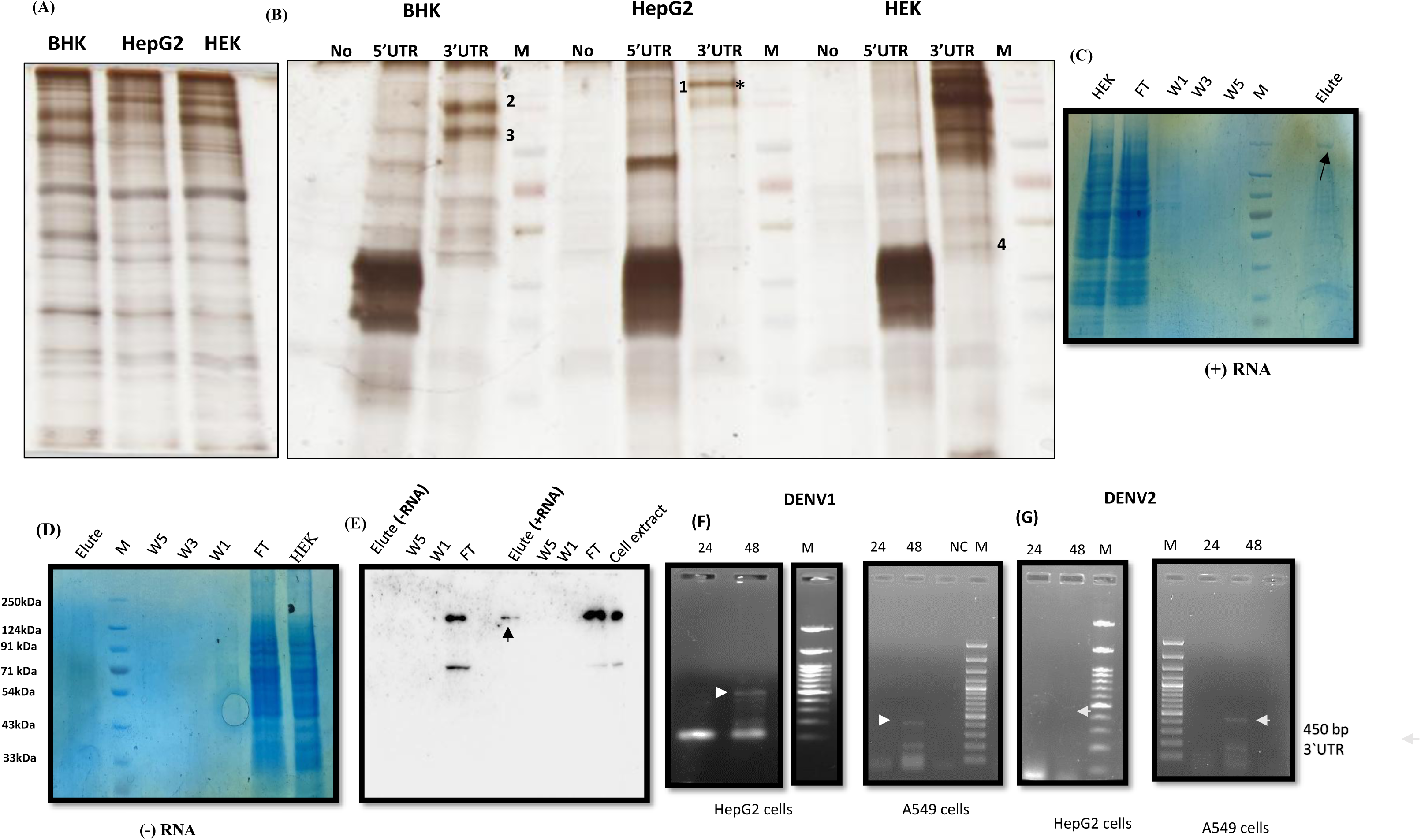
Identification of host proteins that interact with dengue virus 3’UTR. (A) Silver stained 10% SDS PAGE depicting the protein profiles of cytoplasmic extracts from BHK, HepG2, and HEK cell lines. (B) Silver stained 10% SDS PAGE depicting the protein profiles of cytoplasmic extracts that interacted with labelled 5’ and 3’ UTR RNAs. In the negative control, no RNA was included. Interested protein bands that interact with 3’UTR are indicated by numbers and MYH9 as *. (C) & (D) Coomassie stained 10%SDS-PAGE analysis of samples from pulldown assay. A band around 230kDa indicated by an arrow. FT: Flow through, W-Washes, M: Protein Marker. (E) Confirmation of the pulldown experimental data by western blot analysis using MYH9 antibody. (F)&(G) RT-PCR of RNA from immunoprecipitates from DENV1 & DENV2-infected HepG2 & A549 cells. Arrow indicates expected PCR product size of 450 bp. RNA amplified from cells harvested at 24 and 48 h post-infection.

### MYH9 protein levels increase during dengue virus infection

Since MYH9 interacts directly with viral 3’UTR RNA, we hypothesised that it may have additional functions during infections, besides its normal physiological role. In this direction, upon analysing the levels of MYH9 by western blotting under dengue virus infections, the data suggested increased levels of MYH9 in the infected cells (DENV1&2), compared to the mock [Fig 2, A (upper panel), <u>C</u>]. The expression of NS2BNS3 protein was analysed as a marker for infections [Fig 2, A (lower panel), C].

**Figure 2:**
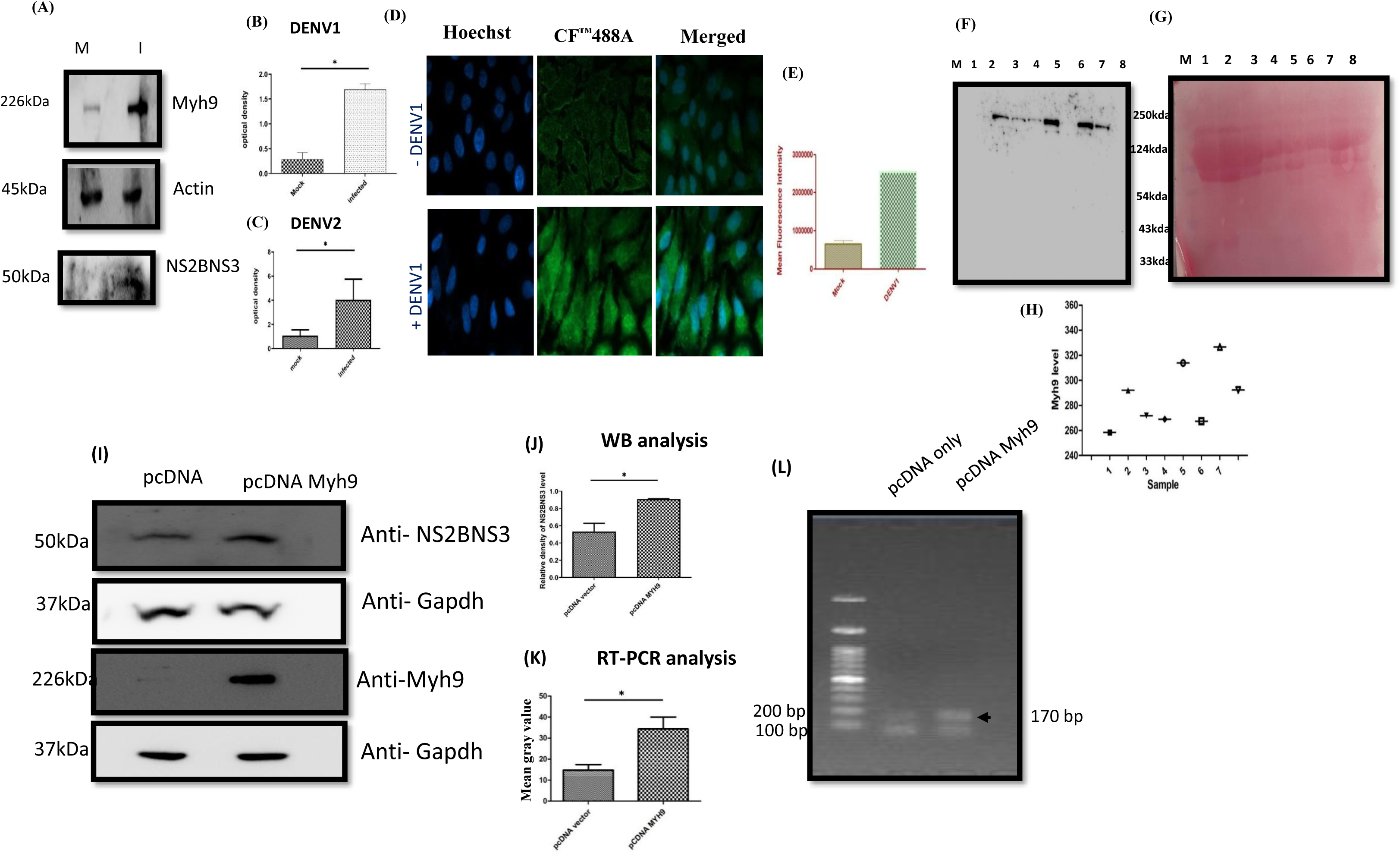
Analysis of the dynamics of MYH9 expression under infection and viral load under MYH9 overexpression. (A) Representative picture of western blot evaluation for MYH9 expression under DENV1&2 infected conditions. The viral protein NS2BNS3 was analysed to indicate the infection. (B) & (C) Bar graph represents the fold increase in expression of MYH9 in DENV1 & DENV2 infected cells. (D) Immunofluorescence image representing MYH9 protein expression: The A549 cells were infected with DENV1 and incubated at 37℃ for 3 days. The cells were then probed using MYH9 antibody and CF^™^488A mouse anti-rabbit IgG after permeabilization. MYH9 is green, and nuclei are blue. (E) The bar graph represents mean fluorescence intensity for mock and infected samples; p<0.002. (F) Representative western blot analysis of MYH9 levels in the dengue virus infected serum samples. (G) Ponceau stained membrane of the above developed blot. (H) Quantitative representation of the bands in the western blot. (1-Healthy, 2-4 DF; 5&6-DHF and 7&8-DSS). (I) Western blot analysis of NS2BNS3 expression in pcDNA MYH9 overexpressed conditions (top panel). Lower panel represent the MYH9 expression. (J) The bar graph to show fold increase in viral load (NS2BNS3) in MYH9 overexpressed cells. (K) Bar diagram representing the fold increase in viral load determined by RT-PCR of culture supernatants with dengue specific primers. (L) Representative agarose gel for the RT-PCR amplified products.

Further, an immunofluorescence assay (IFA) was performed using anti-MYH9 antibodies to verify the above observations. Total MYH9 protein expression was found to be increased after infection when compared with mock cells [Figs 2, D&E]. MYH9 levels were also analysed in the dengue virus infected serum samples. The data indicate that the levels of MYH9 were higher in dengue haemorrhagic fever (DHF) and dengue shock syndrome (DSS) samples compared to the simple dengue fever (DF) and controls [Figs 2, F-H]. All the above studies highlighted increased MYH9 expression during dengue virus infections.

### Overexpression of MYH9 enhances the viral load

Given the elevated MYH9 levels observed during dengue virus infections, we analysed the effect of MYH9 on virus load. In this regard, the protein was overexpressed by transfecting K562 cells with pcDNA-MYH9 and then infecting them with DENV1. The western blot data indicated a significant increase in the viral load under the exogenous expression of MYH9 as compared to the pcDNA vector alone [Figure 2, I-J]. Virus RNA was isolated from the above cell culture supernatants and RT-PCR was analysed that suggested an increase in viral load [Figs 2, K-L]. Both above observations suggest a positive correlation between MYH9 levels and viral load, suggesting a complementary support for each other ie between MYH9 and the virus.

### MYH9 is involved in virus replication

Since MYH9 was found to interact with 3’ UTRs and UTRs are known to be involved in the viral replication process, we presumed its role in the replication of the dengue virus. In this direction, the recruitment of MYH9 to the replication complex on the endoplasmic reticulum (ER) was analysed by double immunofluorescence assay with calnexin (red), an ER marker. The data suggest the localisation of MYH9 (green) on the ER of virus infected cells and was found to be concentrated outside the nucleus [Fig 3, A; lanes vi-x, indicated by arrow]. The fluorescent intensity plots of the above experiment indicated the merg of green (MYH9) and red (ER) colored peaks, supporting the above observations [Fig 3, B; lane ii]. To investigate further in this direction, a co-localization assay was conducted to examine the spatial association between MYH9 and viral double-stranded RNA (dsRNA). The data indicate that dsRNA antibodies (red) bind to the replicative form of the viral RNA that exists in the replication complex of the virus, the location at which the MYH9 (green) was also observed [Fig 3, C; lanes vii-ix]. The fluorescent intensity plot also supported the above observation as the MYH9 expression peak (green) merge with dsRNA expression (red), thus confirming the association of MYH9 with dsRNA suggesting its role virus replication [Fig 3, D; lane ii].

**Figure 3:**
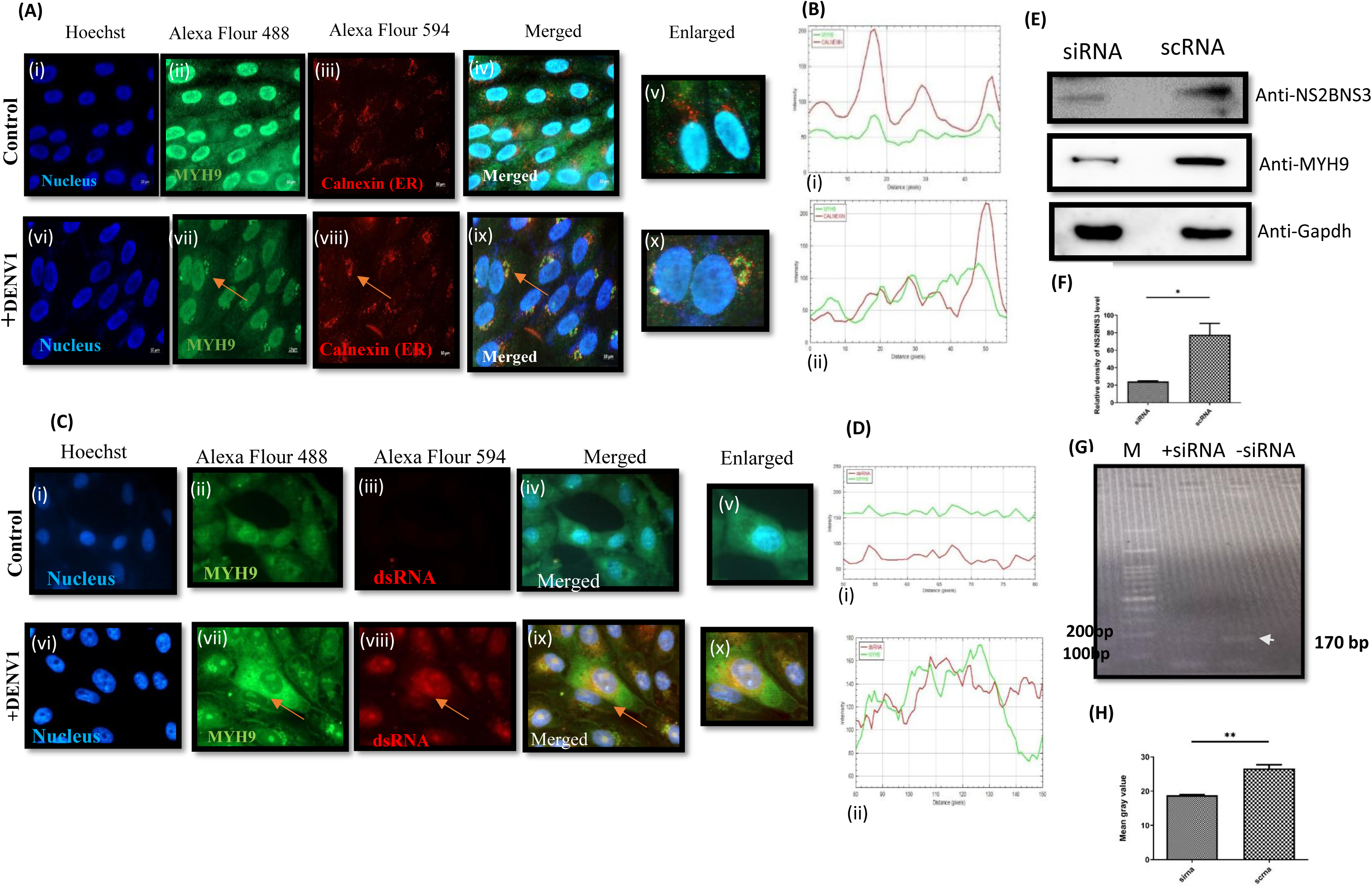
Role of MYH9 in dengue virus genome replication. (A) Co-localization assay of MYH9 with Calnexin in A549 cells after dengue virus infection. Images (i-v) uninfected cells, (vi-ix) dengue infected cells (Arrows represent the MYH9 and calnexin, (x) Enlarged (B) (i) Represents fluorescence intensity plot of uninfected cells showing unmerged green (MYH9) and red (calnexin) colours, (ii) Merged green (MYH9) and red (calnexin) colours in DENV infected cells. (C) Co-localization assay of MYH9 and dsRNA in A549 cells after dengue virus infection. Images (i-v) Represent uninfected cells, (vi-x) Dengue infected cells (Arrows represent the MYH9 and dsRNA, (x) Enlarged. (D) (i) Represents fluorescence intensity plot of uninfected cells showing unmerged green (MYH9) and red (dsRNA) colours, (ii) Merged green (MYH9) and red (dsRNA) colours in DENV infected cells. (E) Representative western blot analysis of dengue virus protein NS2BNS3 upon silencing of *Myh9* mRNA. (F) The bar graph representation of the above experiment. (G) Representative gel picture for RT-PCR analysis of dengue genome (5’end) using the cell culture supernatants under the anti-MYH9 siRNA expression. (H) Bar graph representation of RT-PCR of dengue virus gene segment in siRNA transfected cell supernatant.

### siRNA mediated MYH9 silencing leads to reduced virus load

To validate the involvement of MYH9 in virus replication, we silenced *Myh9* mRNA expression using its siRNA. In this direction, the siRNA was designed, synthesised against MYH9 mRNA and optimised for the concentration that efficiently silences the expression [Fig S7]. After 12-14 hours of incubation after transfecting with siRNA, we found that the virus load was reduced significantly under silencing conditions of MYH9 [Figs 3, E&F]. The reduced levels of virus loads were further confirmed by RT-PCR of the gene segment of the virus using the RNA isolated from the supernatants of the above experiment [Fig 3, G&H].

### MYH9 facilitates dengue virus cellular entry

Literature suggests that the protein MYH9 is involved in the cellular entry of other viruses [18, 19]. Hence, to investigate the role of MYH9 during the early stage of dengue virus infection, Vero-E6 and K562 cells were infected with DENV1 and followed the dynamics of MYH9 at 37°C at 2 and 5 minutes. Interestingly, the data indicated the accumulation of MYH9 at the periphery of the cells after 2 minutes of incubation at 37 °C [Fig 4, A; lanes v-viii]. This observation indicated that MYH9 is involved in dengue virus cellular entry, possibly by acting as a receptor. Since the MYH9 appears to assist the virus entry, and the dengue virus envelope protein is the ligand that interacts with the cellular receptor, we further investigated for the possible MYH9-envelope protein interactions. In this direction, we performed an *in silico* based protein-protein docking assay. The ClusPro docking analysis yielded multiple docking poses, and the lowest energy docked model was selected. The lowest energy model was visualized using PyMOL, revealing a clear interaction interface. The model displayed a strong interaction between the dengue envelope protein and MYH9, with favourable docking energies, suggesting a potential biological interaction between these two proteins [Fig S8] . The interaction surface showed key residues from both proteins forming stable contacts, which may play a role in their potential binding. The visualization highlighted complementary electrostatic and hydrophobic interactions between the dengue envelope protein and MYH9, suggesting that these interactions could be crucial for their functional association. Furthermore, to validate the above observations, a pulldown assay was performed using purified envelope protein and K562 cell extracts. The result showed that a protein band with the molecular weight of approximately 230 kDa was pulled down, which was detected by anti-MYH9 antibodies in western blotting, confirming the envelope-MYH9 interactions [Fig 4, B i&ii &C i&ii]. All the above observations suggest that the protein MYH9 is involved in the cellular entry of the dengue virus by acting as a receptor.

**Figure 4:**
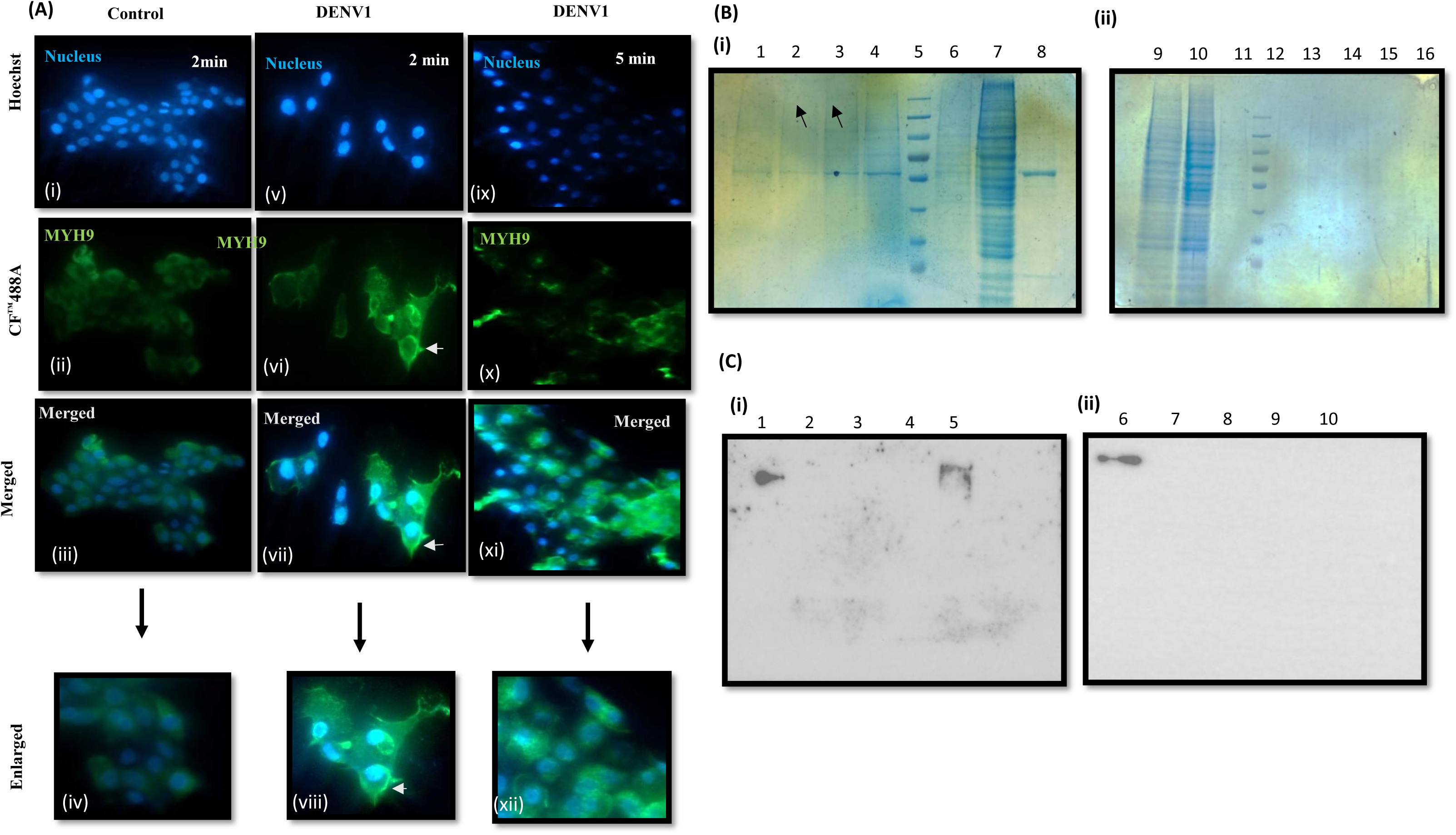
Role of MYH9 in dengue virus cellular entry. (A) Cell-surface expression of MYH9 after dengue viral infection at 4℃, followed by a temperature shift to 37℃. Images (i-iv) Represent uninfected cells, (v-viii) Infected cells after 2 min (surface accumulation is highlighted by arrows) and (ix-xii) Infected after 5 min. (B) Coomassie stained 10% SDS-PAGE analysis for the samples obtained by i*n-vitro* pulldown assay performed using purified envelope protein (i) Represent elutions (E) 2 to 5 (100mM, 150mM, 200mM and 250MM immadazole), M (Marker), wash(W)1 (10mM), Flowthrough (FT) and purified protein in lanes 1-8. (ii). Represents negative control (No envelope protein) SDS gel with lanes 9-16 representing K562 extracts, FT, E2-5, respectively. (C) Western blot confirmation of the above *in vitro* pulldown experiment using MYH9 antibody. i. Represents blot with protein Lanes:1-5 representing FT, W1, E1, 3, 5 ii. Represents samples from negative control, lanes 6-10 FT, W1, E1, 3, 5.

### Anti-MYH9 drug (ML-7) inhibits the dengue virus multiplication

The drug ML-7 is reported to inhibit the MYH9 activity [18, 19]. Hence, we performed indirect immunofluorescence in Vero-E6 cells to examine the inhibitory effect of this drug on dengue virus infection. For this purpose, cells were pretreated with three concentrations of ML-7 (5, 10 & 15 µM) followed by DENV1 infection and an immunofluorescence was carried using the anti-NS2BNS3 antibodies. A significant reduction in fluorescence was observed at 10 and 15µM of the drug concentration [Figs 5 A (xii, xiii, xv &xvi) and B]. RT-PCR analysis was performed using the viral RNA isolated from the supernatants, an amplicon of 170bp (5’ end of the genome) was amplified in the infected sample but the same was absent in the drug treated conditions, confirming the inhibitory role of drug ML-7 [Fig 5, C].

**Figure 5:**
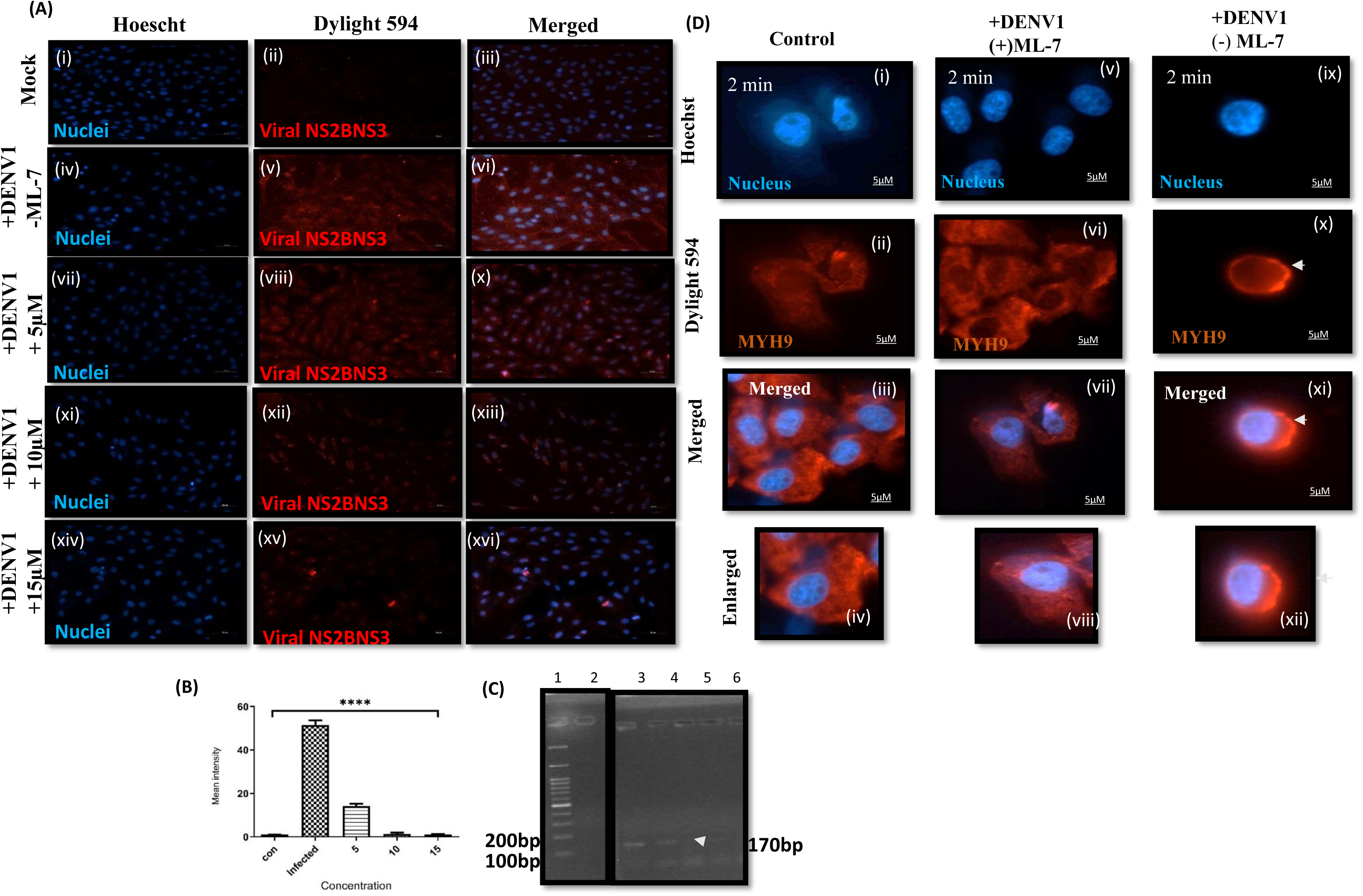
Role of the drug ML-7 on dengue virus replication and cellular entry. (A) Immunofluorescence images representing NS2BNS3 protein expression in Vero cells pre-treated with ML-7 and infected with DENV1. i-iii-Uninfected cells, iv-vii-Infected cells, viii-x: 5µM, xi-xiii: 10µM and xiv-xvi: 15µM of ML-7 treated cells. (B) The bar graph for the above experiment represents total fluorescence intensity for control (No infection), DMSO treated and 5,10,15µM of ML-7 treated cells. (C) Agarose gel electrophoresis gel image of RT-PCR analysis of the viral supernatants from the above experiment. M-Marker, 2-uninfected, 3-Infected and DMSO treated, 4-6: Infected and treated with 5,10,15µM of ML-7. (D) Cell-surface expression of MYH9 after dengue viral adsorption at 4℃ followed by 37℃. Images i-iv are uninfected cells; v-viii are infected cells plus ML-7 treated, ix-xii are infected but not treated (Arrowhead represents peripheral accumulation of MYH9).

### ML-7 inhibits the peripheral accumulation of MYH9 and regains the cellular morphology

It was observed that MYH9 accumulated at the periphery of infected cells during dengue virus infection [Fig 4, A vi, vii&viii]. In order to verify the effect of ML-7 on the above observation, the viral binding and entry assay was performed in the presence of the inhibitor at 15µM concentration. It was found that the accumulation of MYH9 was inhibited under the drug treated conditions [Fig 5, D v-viii]. To analyse the cell morphology of the infected cells after ML-7 treatment, Vero-E6 cells were treated with 5, 10 & 15 µM concentrations of ML-7 followed by DENV1 infection. After 72 hours, the cells exhibited noticeable morphological changes, including a wrinkled appearance with cytopathic effects. But, upon treatment with the drug ML-7, the cell shape was found to changed to normal morphology, and there was no cytopathic effect, suggesting the revival of the cells after drug treatment [Figure S9 ix&x)].

## Discussion

Viruses are etiologic agents of infectious diseases that affect the host system, alter the host’s machinery to favour their multiplication and ultimately disease progression. To do so, viruses interact with many host factors or proteins at every step of life cycle, ie from entry through polyprotein processing, replication, assembly and maturation [16]. Identifying the role of proteins that are involved in the virus life cycle will help in understanding the mechanistic approach of viral genome interactions with host factors that enhance the disease severity. There is limited knowledge of host interacting partners that play a significant role in dengue virus pathogenesis. The 5’ and 3’ untranslated regions (UTRs) of dengue virus are reported to interact with each other as well as host factors [22, 23]. There are proteins reported to interact with dengue virus 3’UTR in both mammalian and mosquito cell lines and deciphering their role in pathogenesis. Proteins like La, IGF-II are reported to bind to the 5’UTR of HCV; others (YB-1, NF-90, DDX-6, PABP) have been implicated in interacting with different regions of 3’UTR and play a role in virus multiplication [24–28]. In this study, to find out the proteins that interact with the dengue virus UTRs, apart from the above, RNA pulldown assay was performed using the biotinylated UDP labelled 5’ and 3’ UTR RNAs and lysates of different cell lines. The analysis indicated many protein bands, indicating the host protein-virus UTR interactions [Fig 1A and B]. Although there were many protein bands for both 5’ and 3’UTRs, this study focused on the proteins that interacted with 3’ UTRs, as this UTR is reported to be more crucial in genome replication. In this direction, a protein band of approximately 230 kDa interacted was eluted and identified. It was found to be the protein MYH9 [Fig 1B; Fig S1A&B], a cytosolic myosin protein that plays an important role in normal cellular processes. Upon doing the western blotting using the eluates of the above experiment with the anti-MYH9 antibodies, the pulled down protein band was confirmed to be the MYH9 [Figs 1, C-E]. The data was further validated in the DENV1&2 infected conditions by RNA immunoprecipitation using the anti-MYH9 antibodies, in which the 3 ’UTRs of both serotypes were amplified from the immunoprecipitants confirming the UTR-MYH9 interactions [Figs 1, F and G].

MYH9 (coded by the gene *Myh9*), a non-muscle myosin protein also termed as non-muscle myosin heavy chain II A belongs to the class II myosin family of which the gene is located on chromosome 22. This family of proteins has different physiological roles, such as cell migration, adhesion, cell shape maintenance, and signal transduction. It is a hexameric protein with two heavy chains, two regulatory light chains and two essential light chains. The light chains control myosin activity and stabilise the heavy chain [29, 30]. Out of 30 motor proteins, the non-muscle myosin is expressed in most of the eukaryotic cells. This protein has been implicated to play a crucial role in pancreatic necrosis virus as it interacts with outer capsid protein VP2 on the cell surface, involved in nanotube formation in porcine reproductive and respiratory syndrome virus infection, facilitating virus spread, initiates viral entry in thrombocytopenia syndrome virus infection through interaction with envelope glycoprotein and herpes simplex virus [18, 20, 31–33]. In the current study, the data suggest that this protein binds to the 3’UTR of dengue virus. *In-silico* analysis further identified the A4 segment of dengue virus 3’UTR variable region that the protein was found to bind. The A4 region is the most conserved region across the dengue virus serotypes, which interact with many host proteins during virus replication and translation [34]. Further, this protein’s binding was found to be localised to the A4 region of 3’ UTR [Figs S2-5], a highly conserved across the dengue virus serotypes [35]. These observations are first ever, although reports suggest the involvement of this protein in the cellular entry of other viruses.

Since the protein interacted with the viral genome, we speculated deviations in its levels during the infections and hence an additional role(s) in the life cycle of the dengue virus. In this direction, upon evaluating its levels during virus infection, we found that they were significantly elevated [Fig 2, A-E and S6]. Further, the protein was also found to be elevated in the dengue virus infected clinical samples [Fig 2, F-H], which further strengthened the above data obtained in the *ex vivo* conditions. On the other hand, viral load was also found to be increased under the MYH9 overexpressed cells [Fig 2, I-L]. These both observations lead us to conclude that the virus induces MYH9 expression, which in turn support elevating the viral load. Since the 3’UTR has been shown to be involved in genome cyclization during virus replication, we further hypothesized the involvement of MYH9 in genome replication. To investigate the involvement of MYH9 in dengue virus replication, we attempted to detect MYH9 on the endoplasmic reticulum (ER), the site of virus replication. The experiments carried using an ER marker calnexin, suggested the presence of the protein in the ER [Fig 3 A (vii-ix) &B(ii)]. There are reports suggesting that dsRNA is the replicative form of the viral RNA that localises to the replication complex of the dengue virus [36]. Hence, to deepen the analysis, the protein’s localization within the replication complex was investigated. It was found that the protein was co-localized with dsRNA [Figs 3, C (vii-ix) and D (ii)]. Additionally, the role of MYH9 on dengue virus replication was evaluated by using the anti-MYH9 siRNA, and the data supported the above findings [Figs 3, E-H]. All the above experimental outcomes established the MYH9’s involvement in dengue virus genome replication.

MYH9 was shown to be involved in virus entry and enhancing the virus load [17–20]. The investigations of the present study suggested the accumulation at the cell periphery immediately after the infection [Figs 4, A vii-ix]. This protein was also found to interact with the dengue virus envelope protein [Figs 4, B, C and S8]. Since the virus induces the MYH9 expression immediately after the infection, it is possible that this protein is forced to evolve acts as a receptor for dengue virus entry during the early stage of infection. At a later stage, the protein localises to the replication complex and interacts with the viral RNA, facilitating genome cyclisation and replication, hence the protein is compiling a dual role in the life cycle of dengue virus ie entry and replication. In this direction, T-cell immunoglobulin and mucin domain 1 (TIM-1), a mitochondrial protein, is also reported to promote the virus by its dual functions, ie entry and autophagy [37]. On the other side, proteins like Ephrin type-A receptor 3 are reported to be utilised at multiple steps of their life cycles by 18 different viruses of different virus families [38, 39]. These observations may provide an opportunity to develop interventions that target many viruses and/or many functions of a single virus.

There are instances suggesting that host proteins could be targeted to reduce the dengue virus production [40]. Similarly, host kinases were also targeted to control these infections [41]. In this direction, MYH9 has been shown to interact with different viral proteins like the envelope glycoprotein of Herpes simplex virus and SARS-CoV2 virus and capsid of pancreatic necrosis virus and facilitates the virus entry into the cells [18,19]. This protein has also been studied for its role in nanotube formation and facilitating viral spread inside the host system [21]. The present study highlighted the role of MYH9 in the life cycle of dengue virus, hence could be considered as a host target to control the infections caused by this virus. In this regard, the anti-MYH9 siRNA developed in this study efficiently inhibited dengue virus infection [Figs 3, E-H]. Furthermore, the drug ML-7 was shown to inhibit the functions of MYH9, hence we extended our studies to analyse the progress of dengue virus infections in the presence of ML-7. The data clearly suggested an inhibitory role of the drug at both the steps (replication & entry) of the life cycle [Figs 5, A-D; S8]. siRNAs and host directed therapeutics (HDTs) were shown to inhibit the viral infections in many instances [40, 41]. These HDTs were reported to have a broad viral spectrum and a lower risk of emerging resistance [42–46]. Hence, the proposed siRNA and/or the drug ML-7, or their relevant molecules, could be used to target the host protein MYH9 and hence to inhibit dengue virus infections.

## Acknowledgements

The authors acknowledge the funding support from SERB, India [EMR/2016/007845 (2018-21)]. The authors also acknowledge Dr. Kishore Vaddadi for the initial experiments carried out in the laboratory for his Ph D thesis.

## Legends for supplementary figures

**Figure S1:** Details of identified 3’UTR-interacting proteins. A. Mascot score profile of host proteins associated with 3′UTR binding. The numbers indicate the spot numbers given on the main Fig. 1A. B. Identified proteins and their functions.

**Figure S2:** *In-silico* prediction of MYH9 binding site on 3’UTR regions. RF and SVM scores in RPIseq analysis using different regions of dengue virus 3’UTR, a score of more than 0.5 was shown in both RF and SVM scores for the A4 region.

**Figure S3.** 3’UTR sequences of dengue virus serotypes. Accession numbers were given for each serotype. The MYH9 binding region was highlighted in red for each serotype.

**Figure S4.** Individual alignment of the A4 regions of 3’UTRs. The 3’UTR of DENV1 with DENV 2, 3 and 4 were presented. The percent identities were indicated by red arrows. The identical nucleotides are represented in stars.

**Figure S5:** Multiple sequence alignment of A4 region of DENV1-4 serotypes using Clustal W, showing conserved nucleotide sequences in the region for the serotypes.

**Figure S6:** RT-PCR analysis of Vero-E6 cell culture supernatants infected with. (A) DENV1 and (B) DENV2 by amplification of the viral NS5 gene fragment showing 493 bp in the 48 hours infected samples. C24/48: control 24/48 hours; I24/48: Infected 24/48 hours

**Figure S7:** Graphical representation of siRNA-mediated knockdown of MYH9. (A) Optimization of siRNA concentration for efficient knockdown of the gene at 50 nM, 100 nM, and 200 nM. Anti-MYH9 antibodies were used to detect the MYH9 levels. (B) Graphical representation of MYH9 levels under different concentrations of siRNA. (C) Graphical representation of Myh9 at 200nM concentration relative to scRNA and NC.

**Figure S8:** Protein-protein docking analysis. A. Crystal structures of dengue virus envelope (4UTC) and MYH9 (4ZME) proteins. The lowest energy docked model was visualized using PyMOL software (The PyMOL Molecular Graphics System Version 3.0.3), showing the interaction interface between the envelope protein and MYH9. B. Table representing possible molecular interactions and weighted scores.

**Figure S9:** Cell morphology analysis of Vero cells under DENV1 infected condition before and after ML-7 treatment. i Represents uninfected, ii-Infected, iii-v-5,10,15 µM of ML-7 treated cells. vi-x Enlarged view of the same cells. Blue arrow indicates CPE:

