## Supplementary figures for "MYH9, a cytosolic myosin protein, binds to dengue virus 3’UTR and facilitates replication and cellular entry"

### Slide 1
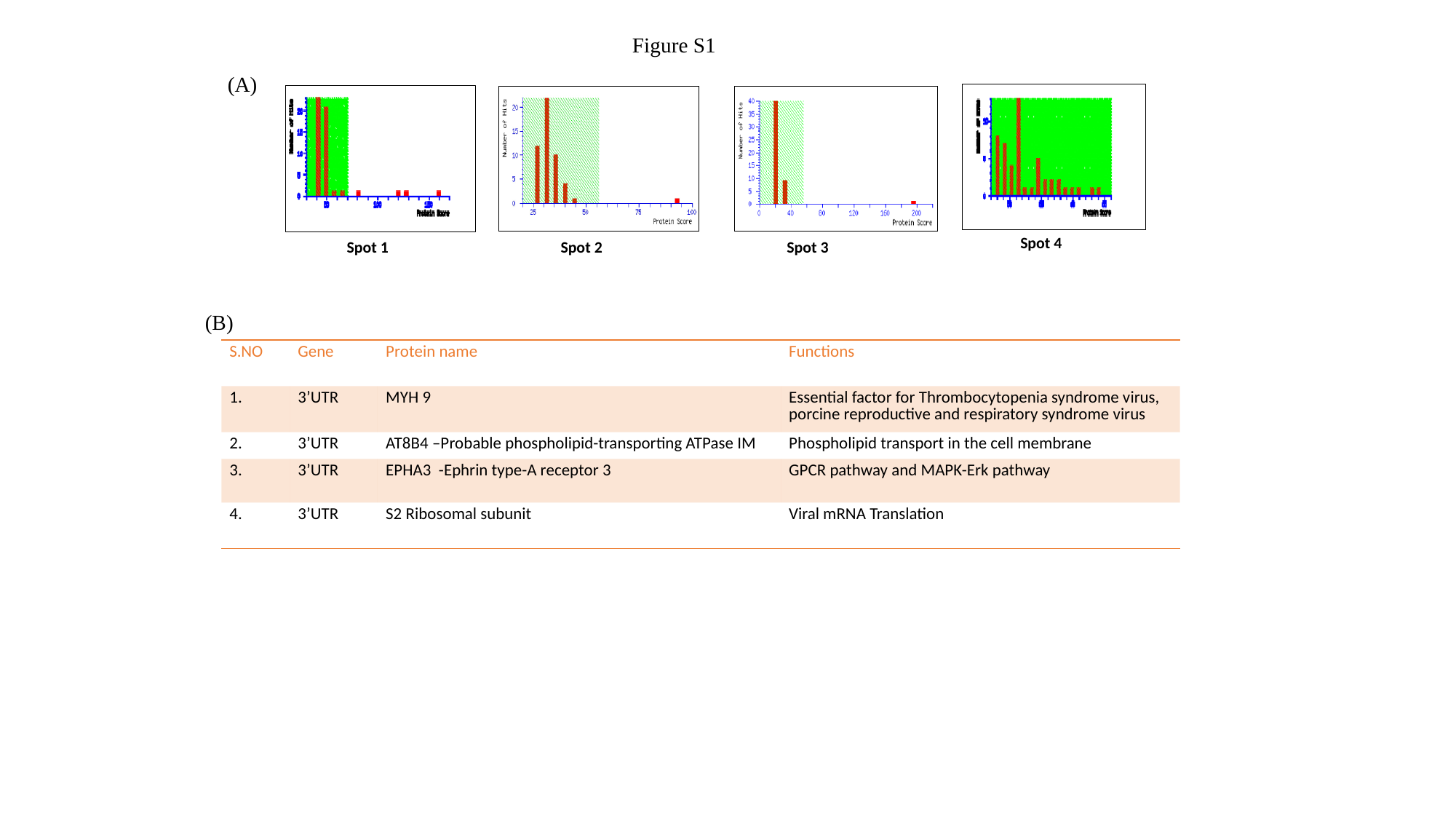

Figure S1
(A)
Spot 4
Spot 1
Spot 2
Spot 3
(B)
| S.NO | Gene | Protein name | Functions |
| --- | --- | --- | --- |
| 1. | 3’UTR | MYH 9 | Essential factor for Thrombocytopenia syndrome virus, porcine reproductive and respiratory syndrome virus |
| 2. | 3’UTR | AT8B4 –Probable phospholipid-transporting ATPase IM | Phospholipid transport in the cell membrane |
| 3. | 3’UTR | EPHA3 -Ephrin type-A receptor 3 | GPCR pathway and MAPK-Erk pathway |
| 4. | 3’UTR | S2 Ribosomal subunit | Viral mRNA Translation |

### Slide 2
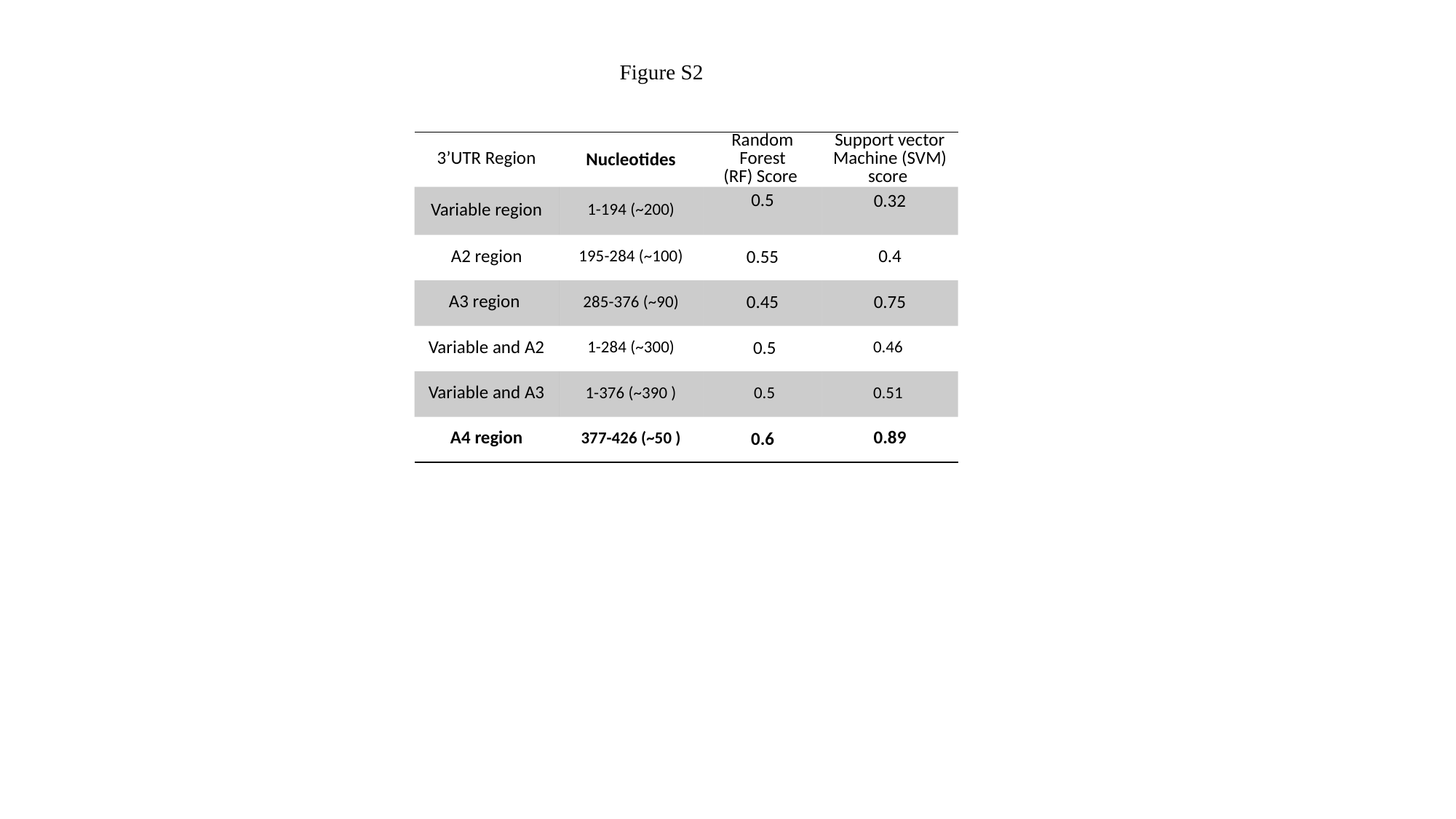

Figure S2
| 3’UTR Region | Nucleotides | Random Forest (RF) Score | Support vector Machine (SVM) score |
| --- | --- | --- | --- |
| Variable region | 1-194 (~200) | 0.5 | 0.32 |
| A2 region | 195-284 (~100) | 0.55 | 0.4 |
| A3 region | 285-376 (~90) | 0.45 | 0.75 |
| Variable and A2 | 1-284 (~300) | 0.5 | 0.46 |
| Variable and A3 | 1-376 (~390 ) | 0.5 | 0.51 |
| A4 region | 377-426 (~50 ) | 0.6 | 0.89 |

### Slide 3
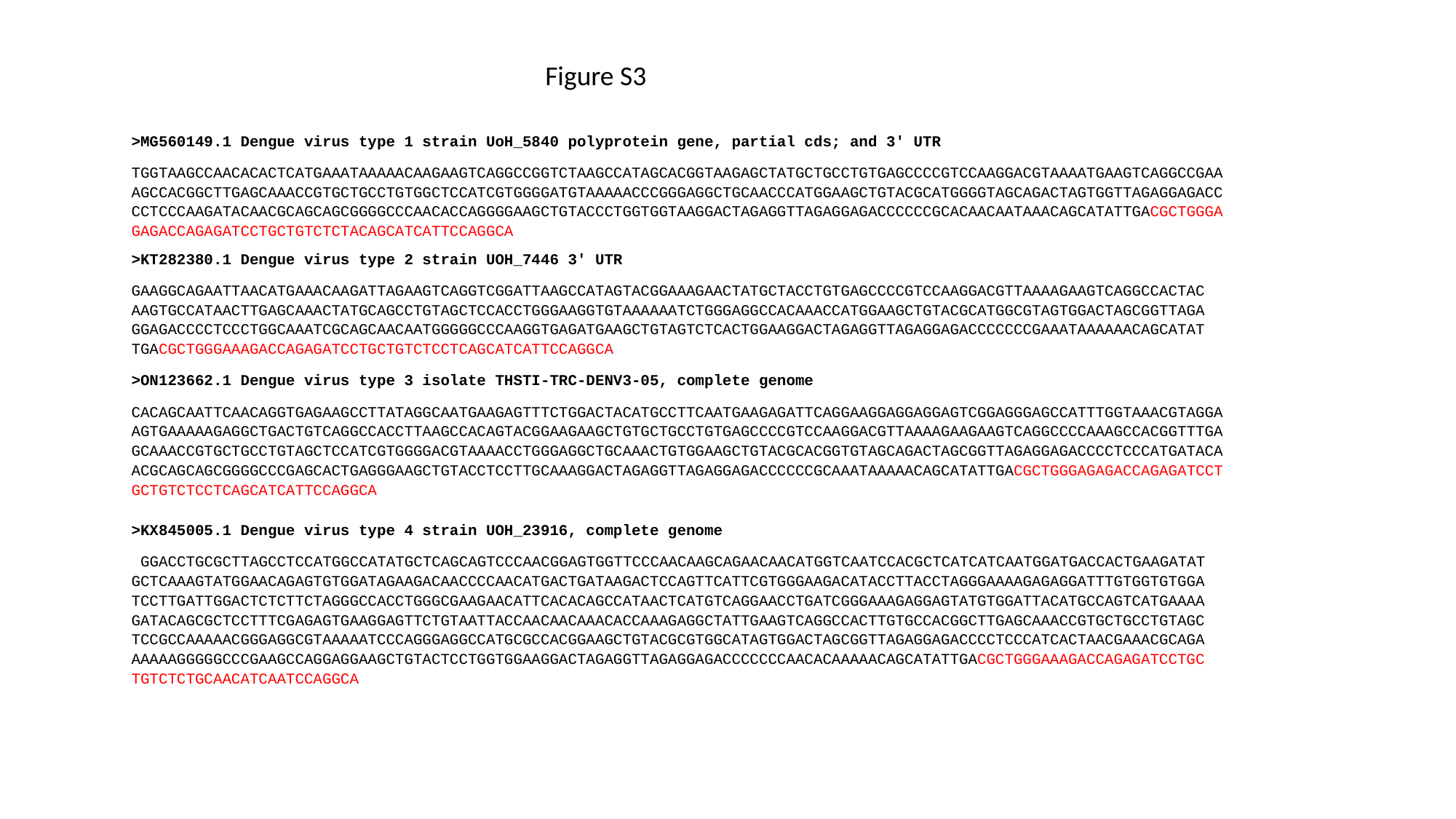

Figure S3
>MG560149.1 Dengue virus type 1 strain UoH_5840 polyprotein gene, partial cds; and 3' UTR
TGGTAAGCCAACACACTCATGAAATAAAAACAAGAAGTCAGGCCGGTCTAAGCCATAGCACGGTAAGAGCTATGCTGCCTGTGAGCCCCGTCCAAGGACGTAAAATGAAGTCAGGCCGAAAGCCACGGCTTGAGCAAACCGTGCTGCCTGTGGCTCCATCGTGGGGATGTAAAAACCCGGGAGGCTGCAACCCATGGAAGCTGTACGCATGGGGTAGCAGACTAGTGGTTAGAGGAGACCCCTCCCAAGATACAACGCAGCAGCGGGGCCCAACACCAGGGGAAGCTGTACCCTGGTGGTAAGGACTAGAGGTTAGAGGAGACCCCCCGCACAACAATAAACAGCATATTGACGCTGGGAGAGACCAGAGATCCTGCTGTCTCTACAGCATCATTCCAGGCA
>KT282380.1 Dengue virus type 2 strain UOH_7446 3' UTR
GAAGGCAGAATTAACATGAAACAAGATTAGAAGTCAGGTCGGATTAAGCCATAGTACGGAAAGAACTATGCTACCTGTGAGCCCCGTCCAAGGACGTTAAAAGAAGTCAGGCCACTACAAGTGCCATAACTTGAGCAAACTATGCAGCCTGTAGCTCCACCTGGGAAGGTGTAAAAAATCTGGGAGGCCACAAACCATGGAAGCTGTACGCATGGCGTAGTGGACTAGCGGTTAGAGGAGACCCCTCCCTGGCAAATCGCAGCAACAATGGGGGCCCAAGGTGAGATGAAGCTGTAGTCTCACTGGAAGGACTAGAGGTTAGAGGAGACCCCCCCGAAATAAAAAACAGCATATTGACGCTGGGAAAGACCAGAGATCCTGCTGTCTCCTCAGCATCATTCCAGGCA
>ON123662.1 Dengue virus type 3 isolate THSTI-TRC-DENV3-05, complete genome
CACAGCAATTCAACAGGTGAGAAGCCTTATAGGCAATGAAGAGTTTCTGGACTACATGCCTTCAATGAAGAGATTCAGGAAGGAGGAGGAGTCGGAGGGAGCCATTTGGTAAACGTAGGAAGTGAAAAAGAGGCTGACTGTCAGGCCACCTTAAGCCACAGTACGGAAGAAGCTGTGCTGCCTGTGAGCCCCGTCCAAGGACGTTAAAAGAAGAAGTCAGGCCCCAAAGCCACGGTTTGAGCAAACCGTGCTGCCTGTAGCTCCATCGTGGGGACGTAAAACCTGGGAGGCTGCAAACTGTGGAAGCTGTACGCACGGTGTAGCAGACTAGCGGTTAGAGGAGACCCCTCCCATGATACAACGCAGCAGCGGGGCCCGAGCACTGAGGGAAGCTGTACCTCCTTGCAAAGGACTAGAGGTTAGAGGAGACCCCCCGCAAATAAAAACAGCATATTGACGCTGGGAGAGACCAGAGATCCTGCTGTCTCCTCAGCATCATTCCAGGCA
>KX845005.1 Dengue virus type 4 strain UOH_23916, complete genome
 GGACCTGCGCTTAGCCTCCATGGCCATATGCTCAGCAGTCCCAACGGAGTGGTTCCCAACAAGCAGAACAACATGGTCAATCCACGCTCATCATCAATGGATGACCACTGAAGATATGCTCAAAGTATGGAACAGAGTGTGGATAGAAGACAACCCCAACATGACTGATAAGACTCCAGTTCATTCGTGGGAAGACATACCTTACCTAGGGAAAAGAGAGGATTTGTGGTGTGGATCCTTGATTGGACTCTCTTCTAGGGCCACCTGGGCGAAGAACATTCACACAGCCATAACTCATGTCAGGAACCTGATCGGGAAAGAGGAGTATGTGGATTACATGCCAGTCATGAAAAGATACAGCGCTCCTTTCGAGAGTGAAGGAGTTCTGTAATTACCAACAACAAACACCAAAGAGGCTATTGAAGTCAGGCCACTTGTGCCACGGCTTGAGCAAACCGTGCTGCCTGTAGCTCCGCCAAAAACGGGAGGCGTAAAAATCCCAGGGAGGCCATGCGCCACGGAAGCTGTACGCGTGGCATAGTGGACTAGCGGTTAGAGGAGACCCCTCCCATCACTAACGAAACGCAGAAAAAAGGGGGCCCGAAGCCAGGAGGAAGCTGTACTCCTGGTGGAAGGACTAGAGGTTAGAGGAGACCCCCCCAACACAAAAACAGCATATTGACGCTGGGAAAGACCAGAGATCCTGCTGTCTCTGCAACATCAATCCAGGCA

### Slide 4
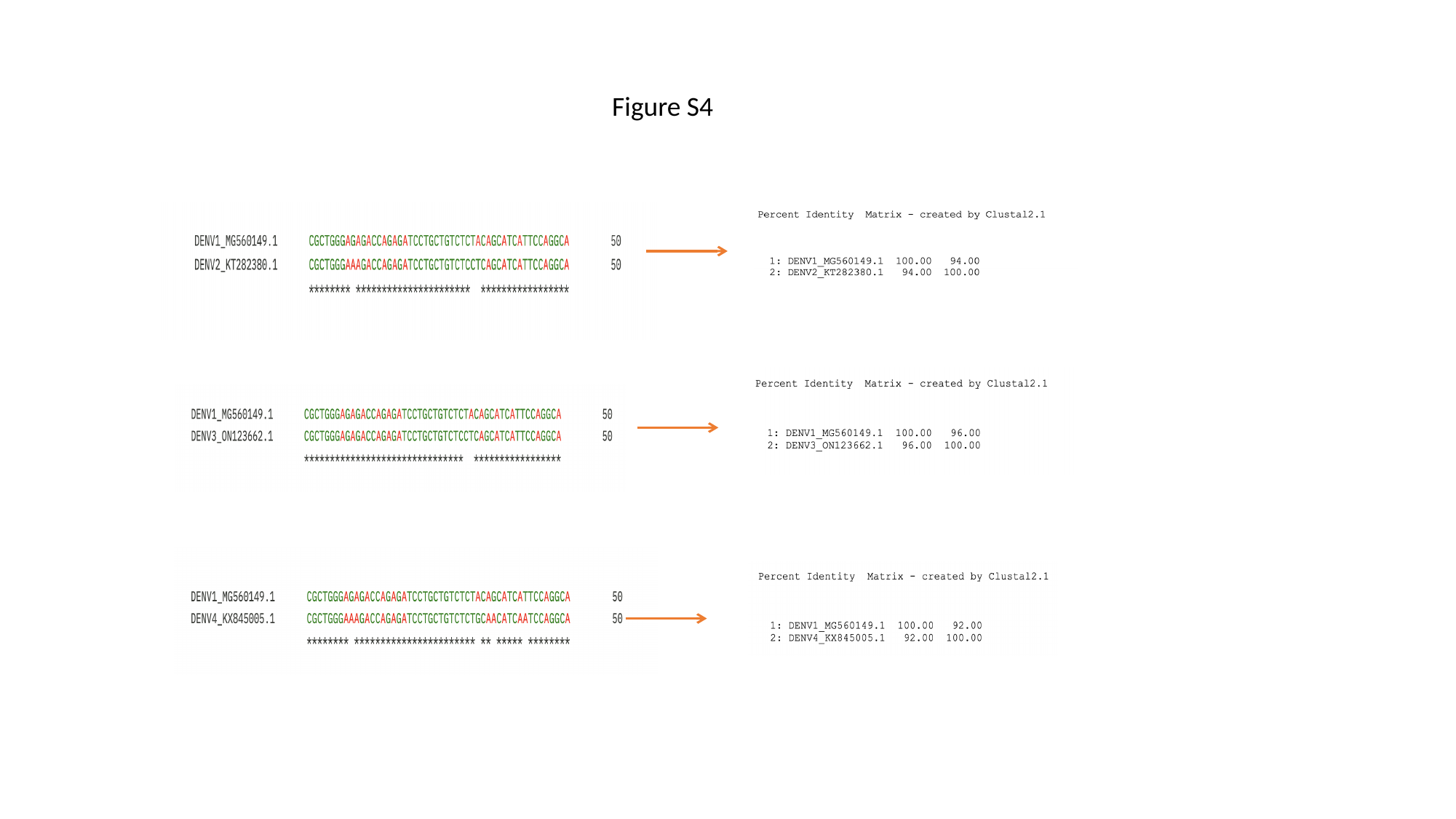

#### Figure S4

### Slide 5
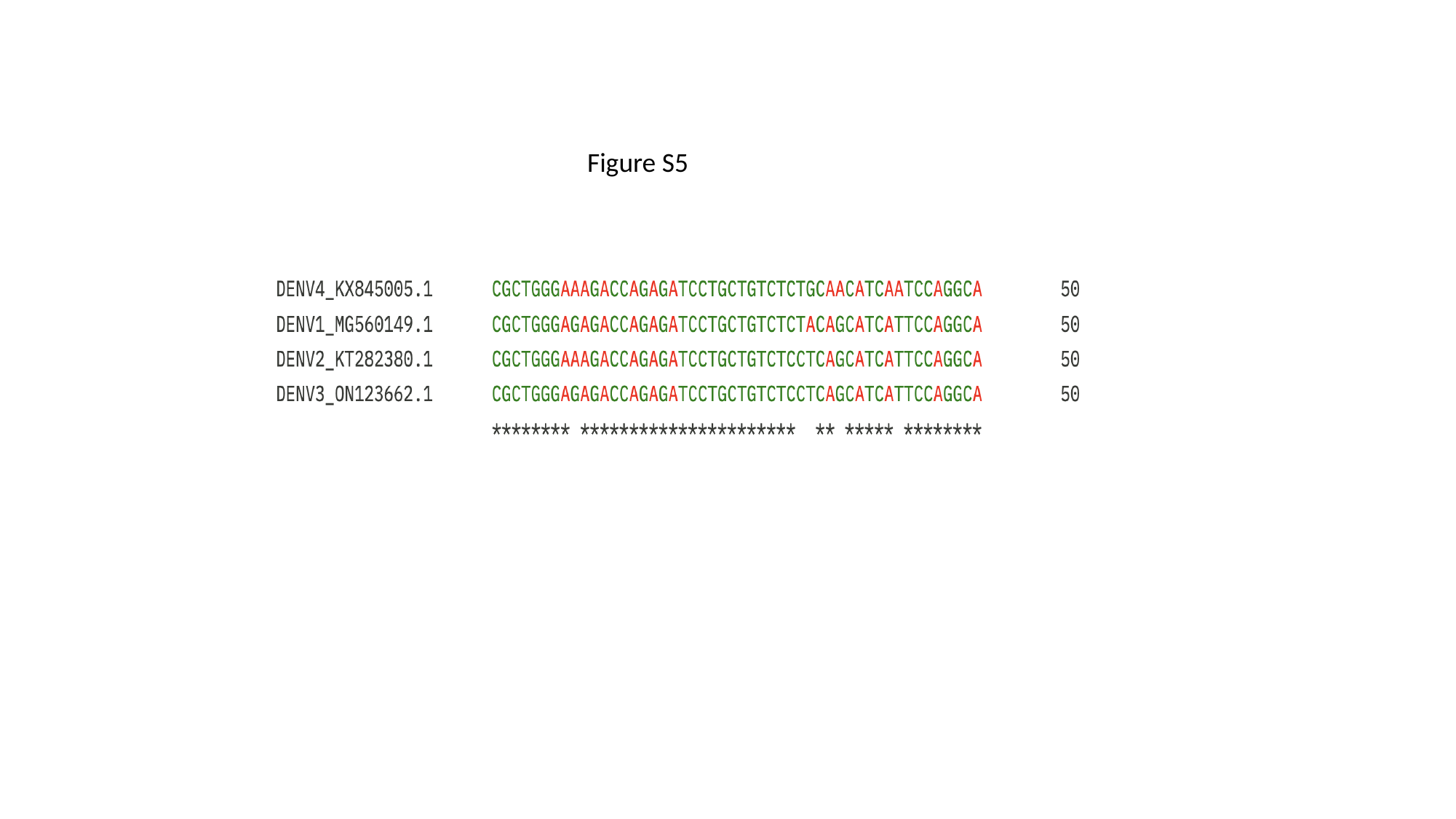

#### Figure S5

### Slide 6
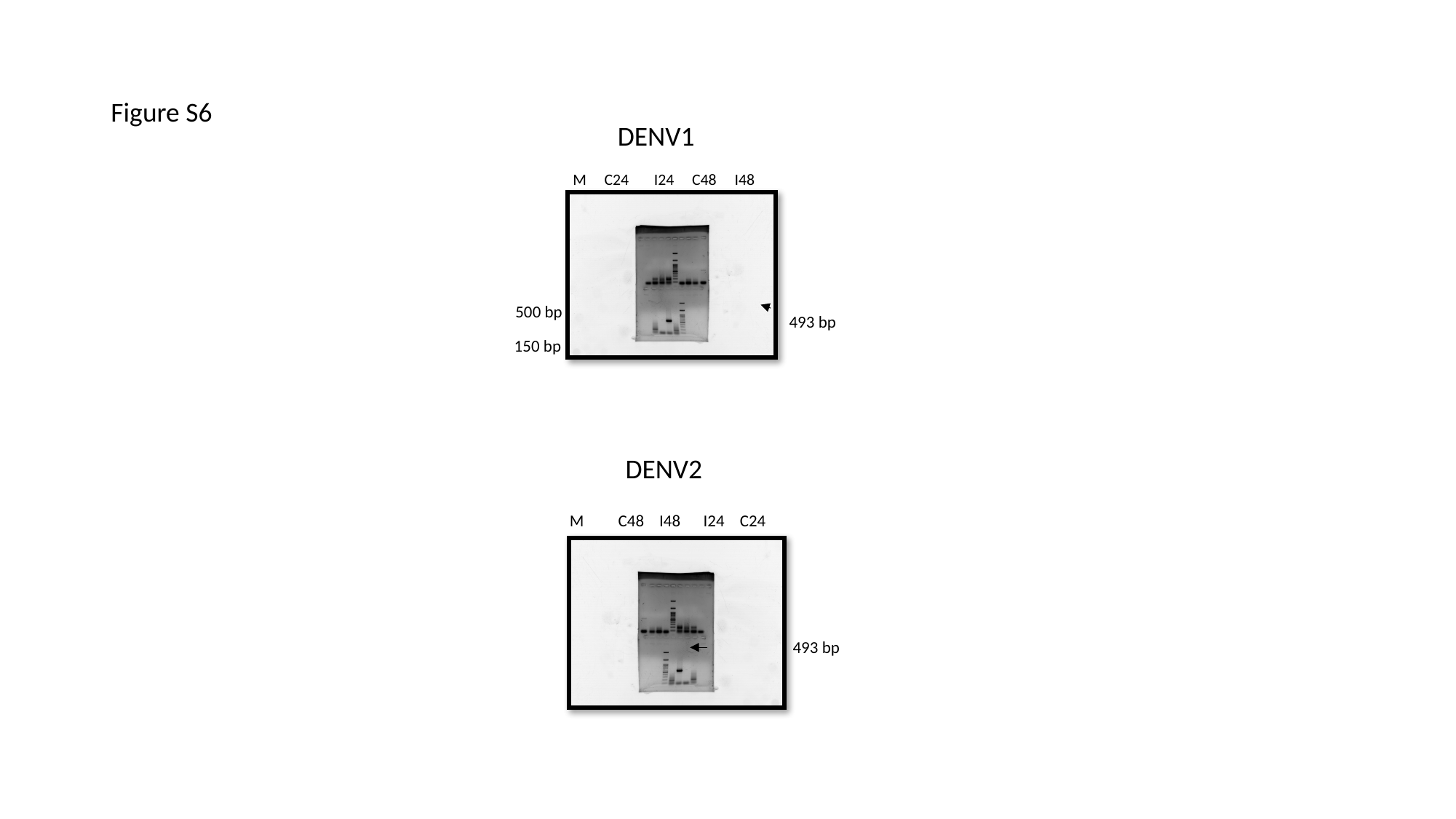

#### Figure S6
DENV1
 M C24 I24 C48 I48
500 bp
150 bp
M C48 I48 I24 C24
493 bp
493 bp
DENV2

### Slide 7
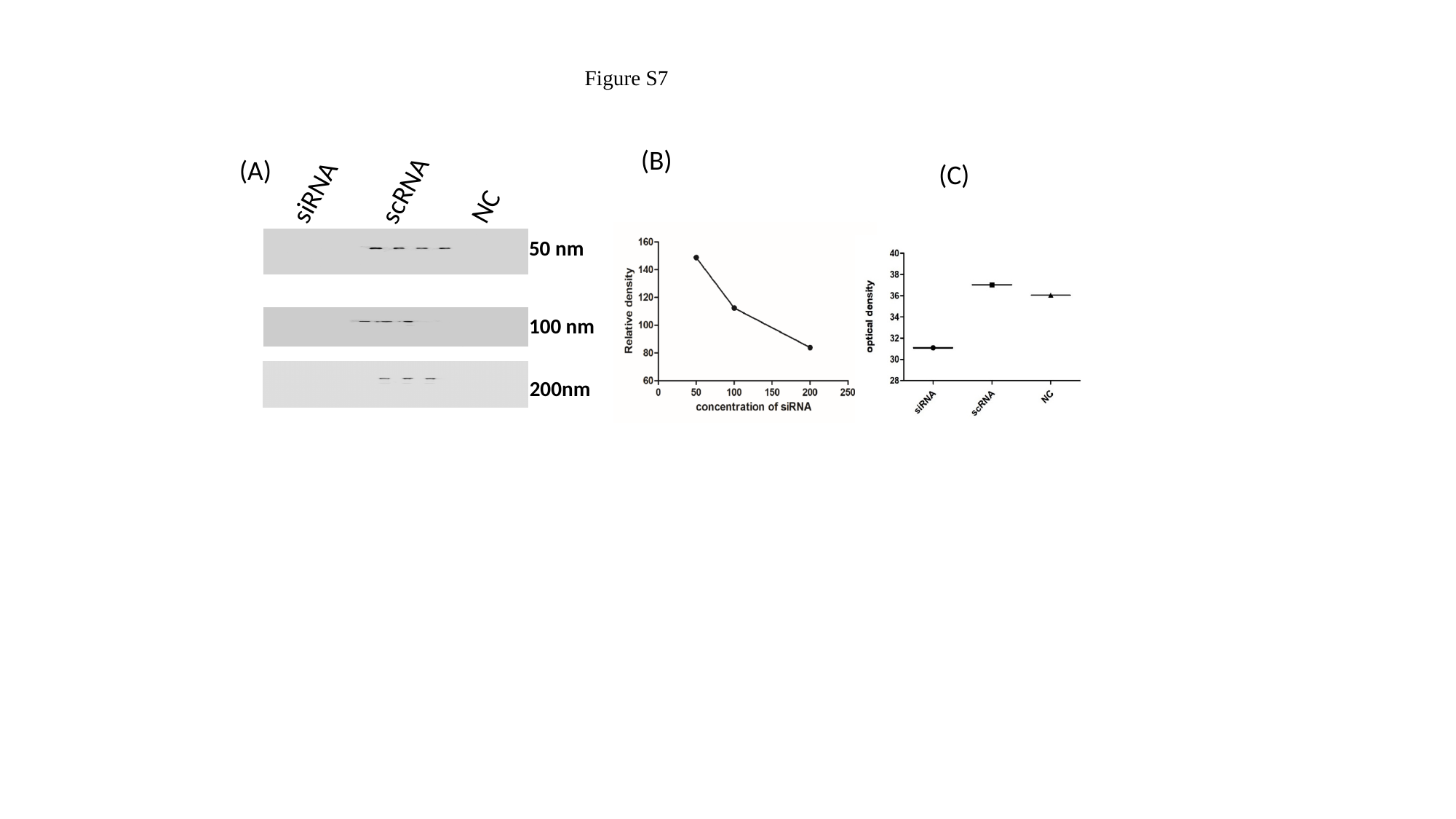

Figure S7
(B)
(A)
(C)
siRNA
scRNA
NC
50 nm
100 nm
200nm

### Slide 8
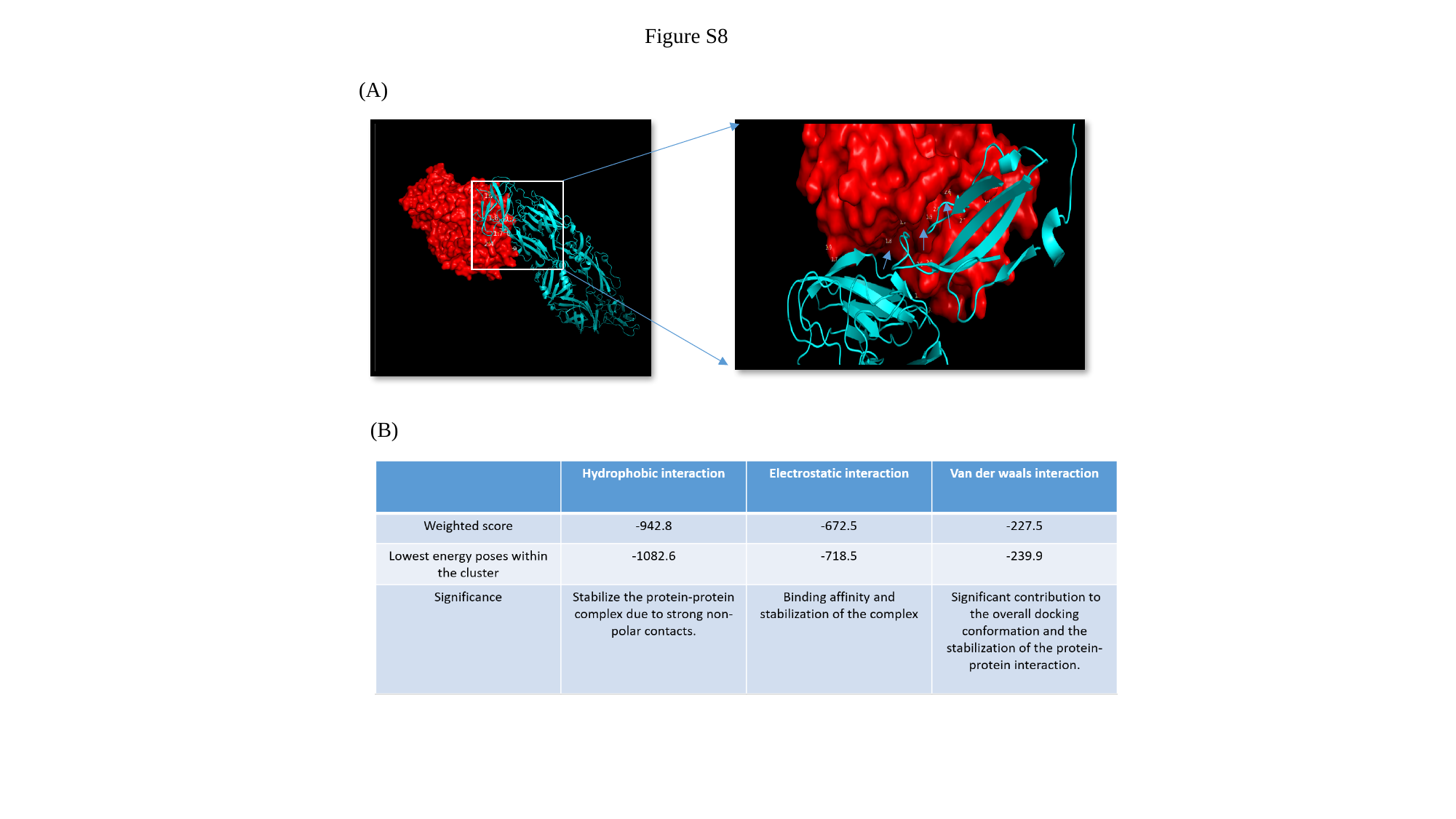

Figure S8
(A)
(B)

### Slide 9
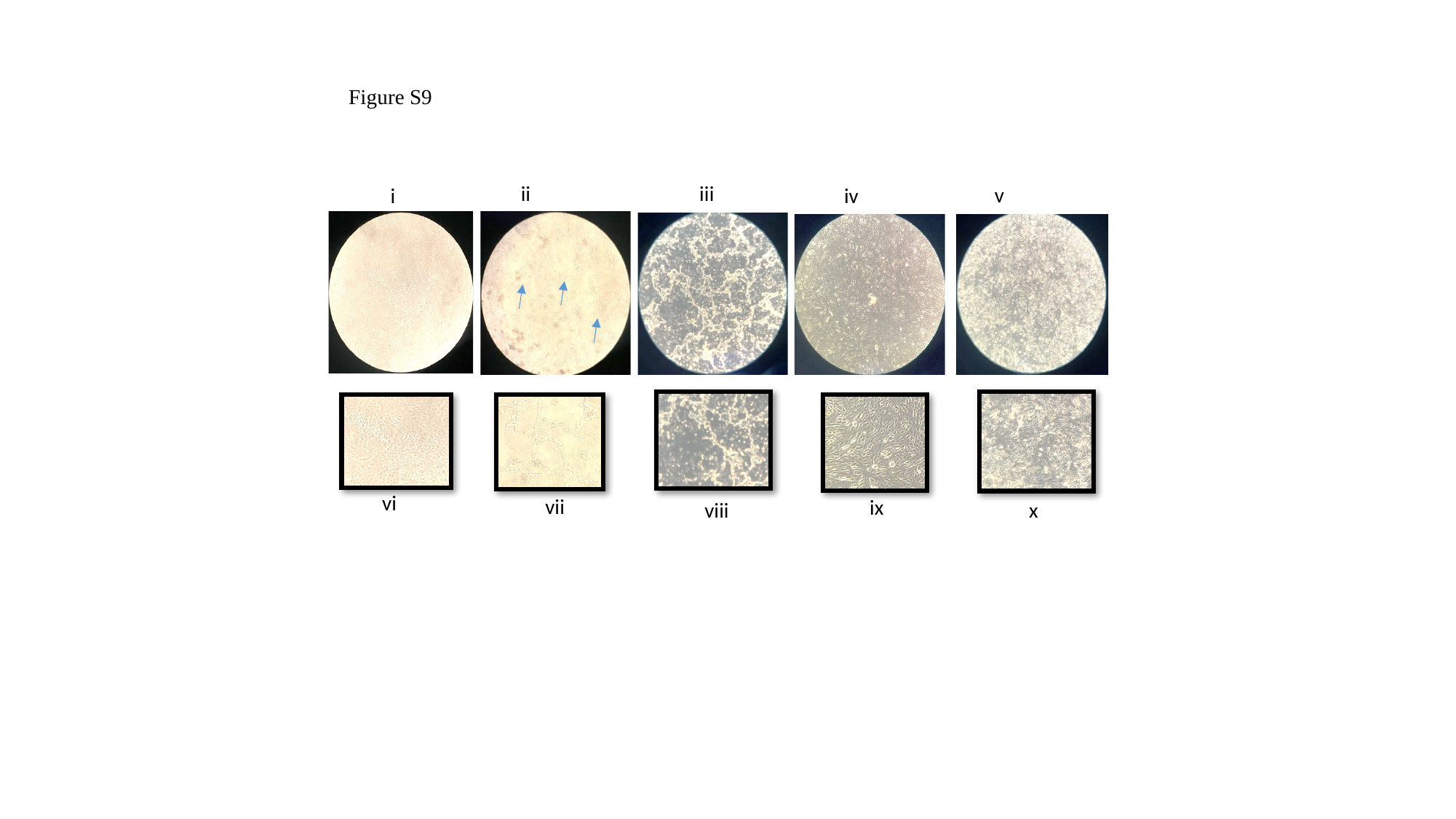

Figure S9
iii
ii
v
iv
i
vi
vii
ix
viii
x
